# Autosomal admixture levels are informative about sex bias in admixed populations

**DOI:** 10.1101/006452

**Authors:** Amy Goldberg, Paul Verdu, Noah A Rosenberg

**Affiliations:** Department of Biology, Stanford University, Stanford, CA, 94305-5020 USA; CNRS-MNHN-Université Paris Diderot, UMR7206 Ecoanthropology and Ethnobiology, Paris, France

## Abstract

Sex-biased admixture has been observed in a wide variety of admixed populations. Genetic variation in sex chromosomes and ratios of quantities computed from sex chromosomes and autosomes have often been examined in order to infer patterns of sex-biased admixture, typically using statistical approaches that do not mechanistically model the complexity of a sex-specific history of admixture. Here, expanding on a model of Verdu & Rosenberg (2011) that did not include sex specificity, we develop a model that mechanistically examines sex-specific admixture histories. Under the model, multiple source populations contribute to an admixed population, potentially with their male and female contributions varying over time. In an admixed population descended from two source groups, we derive the moments of the distribution of the autosomal admixture fraction from a specific source population as a function of sex-specific introgression parameters and time. Considering admixture processes that are constant in time, we demonstrate that surprisingly, although the mean autosomal admixture fraction from a specific source population does not reveal a sex bias in the admixture history, the variance of autosomal admixture is informative about sex bias. Specifically, the long-term variance decreases as the sex bias from a contributing source population increases. This result can be viewed as analogous to the reduction in effective population size for populations with an unequal number of breeding males and females. Our approach can contribute to methods for inference of the history of complex sex-biased admixture processes by enabling consideration of the effect of sex-biased admixture on autosomal DNA.

## Introduction

Populations often experience sex-biased demographic processes, in which males and females contributing to the gene pool of a population are drawn from source groups in different proportions, owing to patterns of inbreeding avoidance, dispersal, and mating practices (Pusey 1987; Lawson Handley & Perrin 2007). In humans, sex-biased demography has had a particular effect on admixed populations, populations that have often been founded or influenced by periods of colonization and forced migration involving an initial or continuing admixture process (Mesa et al. 2000; Seielstad 2000; Wilkins & Marlowe 2006; Tremblay & Vezina 2010; Heyer et al. 2012).

Genetic signatures of sex-biased admixture have been empirically investigated in a variety of human populations. In the Americas, these include African American, Latino, and Native American populations (Bolnick et al. 2006; Wang et al. 2008; Stefflova et al. 2009; Tishkoff et al. 2009; Bryc et al. 2010a,b; Moreno-Estrada et al. 2013; Verdu et al. 2014). Sex-biased admixture and migration have also been examined in populations throughout Asia (Oota et al. 2001; Wen et al. 2004; Chaix et al. 2007; Ségurel et al. 2008; Chaubey et al. 2011; Pemberton et al. 2012; Pijpe et al. 2013), Austronesia (Kayser et al. 2003, 2006, 2008; Cox et al. 2010; Lansing et al. 2011) and Africa (Wood et al. 2005; Tishkoff et al. 2007; Berniell-Lee et al. 2008; Beleza et al. 2013; Petersen et al. 2013; Verdu et al. 2013).

Sex-specific admixture and migration processes have typically been studied using comparisons of the Y chromosome, which is paternally inherited, and the mitochondrial genome, inherited maternally (Seielstad et al. 1998; Oota et al. 2001; Wood et al. 2005; Bolnick et al. 2006; Gunnarsdóttir et al. 2011; Lacan et al. 2011). More recently, as the Y chromosome and mitochondrial genome each represent single nonrecombining loci that provide an incomplete genomic perspective, sex-biased admixture has been examined by comparisons of autosomal DNA to the X chromosome (Lind et al. 2007; Wang et al. 2008; Bryc et al. 2010a,b; Cox et al. 2010; Beleza et al. 2013; Verdu et al. 2013).

The Y–mitochondrial and X–autosomal frameworks are both sensible, as both involve comparisons of two types of loci that follow different modes of inheritance in males and females. What has not been clear, however, is that autosomal data, which have not typically been viewed as the most informative loci for studies of sex-specific processes, can carry information about sex-biased admixture, even in the absence of a comparison with other components of the genome.

We demonstrate this surprising result through an extension of a mechanistic model for the admixture history of a hybrid population. In a diploid autosomal framework, Verdu & Rosenberg (2011) examined contributions of multiple source populations that varied through time, without considering sex specificity. Here, expanding on the model of Verdu & Rosenberg (2011), we develop a model that mechanistically considers sex-specific admixture histories in which multiple source populations contribute to the admixed population, potentially with varying female and male contributions across generations (Fig. 1). In an admixed population descended from two source populations, we derive the moments of the distribution of the fraction of autosomal admixture from a specific source population, as a function of sex-specific admixture parameters and time. We analyze the behavior of the model, considering admixture processes that are constant in time, and we show that the moments contain information about the sex bias.

**Figure 1:**
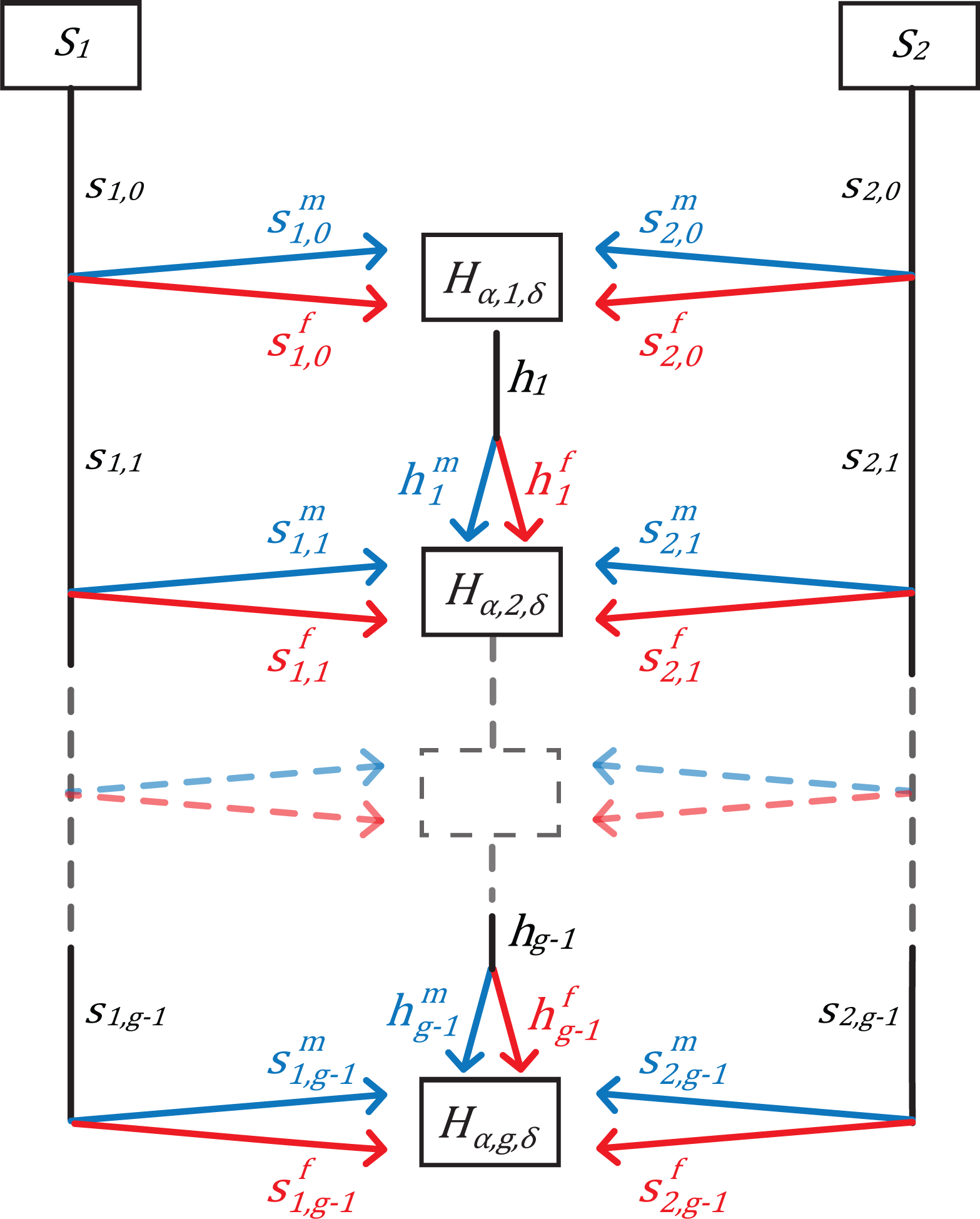
Schematic of the mechanistic model of admixture over time. Two source populations, *S*_1_ and *S*_2_, contribute both males and females to the next generation of the hybrid population *H*, potentially with time-varying proportions. The fractional contributions of the source populations and the hybrid population to the next generation *G* are *s*_1,*g*_, *s*_2,*g*_ and *h_g_*, respectively. Sex-specific contributions from the populations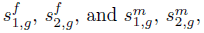 for females and males respectively. *H_α,g,δ_* represents the fraction of admixture from source population *α* ∈ {1, 2*}* in generation *g* for a random individual of sex *δ ∈ {f, m}* in population *H*.

## The model

Several studies have described mechanistic models of admixture (Chakraborty & Weiss 1988; Long 1991; Ewens & Spielman 1995; Guo et al. 2005; Verdu & Rosenberg 2011; Gravel 2012; Jin et al. 2013). We follow the notation and style of the model of Verdu & Rosenberg (2011), studying a hybrid population, *H,* which consists of immigrant individuals from *M* isolated source populations and hybrid individuals who have ancestors from two or more source populations. The source populations are labeled *S_α_*, for *α* from 1 to *M*. We focus on the case of *M* = 2.

We define the parameters *s_α,g−_*_1_ and *h_g−_*_1_ as the contributions from source populations *S_α_* and *H*, respectively, to the gene pool of the hybrid population *H* at the next generation, *g*. That is, for a randomly chosen individual at generation *g*, the probabilities that a randomly chosen parent of the individual derives from *S_α_* and *H* are *s_α,g−_*_1_ and *h_g−_*_1_, respectively. We define the sex-specific parameter 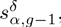 for *δ ∈* {*f, m*}, as the probability that the type-*δ* parent of a randomly chosen individual from the hybrid population at generation *g* is from source population *S_α_*. Similarly, 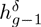 is the probability that the type-*δ* parent of a randomly chosen individual in *H* at generation *g* is from *H* itself. We consider a two-sex model, using *f* for female and *m* for male. Thus, because each individual has one parent of each type, female and male, we have

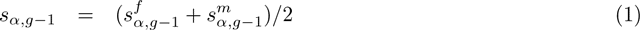

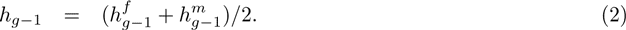

The contributions to the next generation of the three source populations (*S*_1_*, S*_2_*, H*) sum to one:

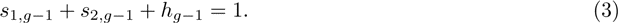

Similarly, the female and male contributions to the next generation separately sum to one,

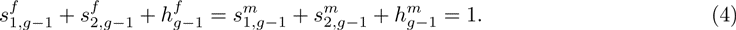

At the first generation, *g* = 1, the hybrid population has not previously existed; therefore,

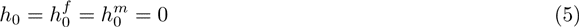

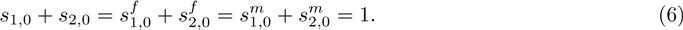

The first generation has two independent parameters, 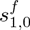 and 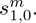 Each subsequent generation contributes four additional independent parameters 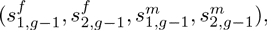 and considering the first *g* generations, there are 4 *g −* 2 independent parameters. The model is discrete in time and assumes non-overlapping generations.

Our model allows us to consider complex sex-biased admixture processes by allowing uneven sex-specific contributions from each source population at each generation. It reduces to the model of Verdu & Rosenberg (2011) when the sex-specific contributions are equal within a source population, that is, if for each *g*, 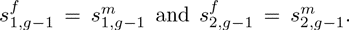 We perform similar computations to those of Verdu & Rosenberg (2011), illustrating that in certain cases, our results reduce to those obtained when sex specificity is not considered.

We let *L* be a random variable indicating the source populations of the parents of a random individual from the hybrid population, *H*. *L* takes its values from the set of all possible ordered parental combinations, {*S*_1_*S*_1_*, S*_1_*H, S*_1_*S*_2_*, HS*_1_*, HH, HS*_2_*, S*_2_*S*_1_*, S*_2_*H, S*_2_*S*_2_}, listing the female parent first. We assume random mating in the hybrid population at each generation, so that the probability that an offspring has a particular pair of source populations for his or her parents is simply the product of the probabilities for having the female and male parents (Table 1).

**Table 1:** The probabilities that an individual from the hybrid population at generation *g* has one of nine possible sets of parents from *S*_1_*, S*_2_ or *H*, assuming random mating. The parameter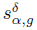 is the probability that the parent of sex *δ* for a randomly chosen individual from the hybrid population, at generation *g*, is from the source population *α*. Similarly, the probability that this parent is from *H* is 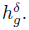

| Case | Female Parent's Population | Male Parent's Population | Probability |
| --- | --- | --- | --- |
| 1 | $S_1$ | $S_1$ | $s_{1,g-1}^f s_{1,g-1}^m$ |
| 2 | $S_1$ | $H$ | $s_{1,g-1}^f h_{g-1}^m$ |
| 3 | $S_1$ | $S_2$ | $s_{1,g-1}^f s_{2,g-1}^m$ |
| 4 | $H$ | $S_1$ | $h_{g-1}^f s_{1,g-1}^m$ |
| 5 | $H$ | $H$ | $h_{g-1}^f h_{g-1}^m$ |
| 6 | $H$ | $S_2$ | $h_{g-1}^f s_{2,g-1}^m$ |
| 7 | $S_2$ | $S_1$ | $s_{2,g-1}^f s_{1,g-1}^m$ |
| 8 | $S_2$ | $H$ | $s_{2,g-1}^f h_{g-1}^m$ |
| 9 | $S_2$ | $S_2$ | $s_{2,g-1}^f s_{2,g-1}^m$ |

We define the fraction of admixture, the random variable *H_α,g,δ_*, as the probability that an autosomal genetic locus in a random individual of sex *δ* from the hybrid population in generation *g* ultimately originates from source population *α*. The sex-specific fractions of admixture are related to the total fraction of admixture *H_α,g_* from source population *α* in generation *g* by *H_α,g_* = (*H_α,g,f_* + *H_α,g,m_*)*/*2.

Under the model, we derive expressions for the moments of the fraction of admixture. Autosomal DNA is inherited non-sex-specifically and from both parents; therefore, female and male offspring have identical distributions of admixture, and *H_α,g,f_* and *H_α,g,m_* are identically distributed. Each of these quantities depends on both the female fraction and the male fraction of admixture in the previous generation, but conditional on the previous generation (that is, on *H_α,g−_*_1_*_,f_* and *H_α,g−_*_1_*_,m_*), they are independent. For our two-population model, we consider the non-sex-specific fraction of admixture, *H*_1_*_,g,δ_*, treating *δ* here as representing either *f* or *m,* (but retaining the same meaning throughout). The quantity *H*_1_*_,g,δ_* depends on both sex-specific fractions of admixture from the previous generation, *H*_1_*_,g−_*_1_*_,f_* and *H*_1_*_,g−_*_1_*_,m_*.

## Distribution of the admixture fraction from a specific source

The definition of the model parameters and the values from Table 1 allow us to write a recursion relation for the fraction of admixture from source population 1 for a random individual of sex *δ* from the hybrid population at generation *g*, or *H*_1_*_,g,δ_*. For the first generation, *g* = 1, we have

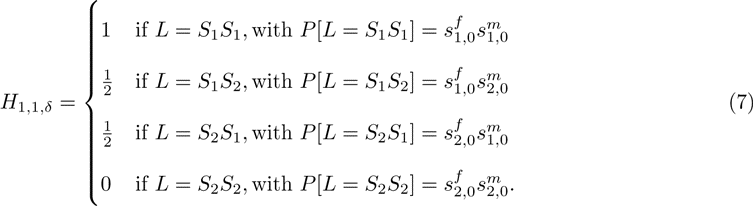

For all subsequent generations, *g ≥* 2, we have

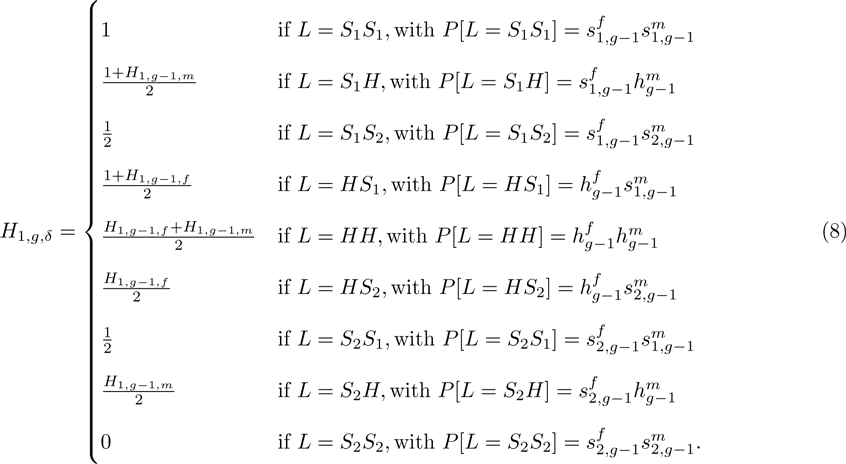

Using eqs. (7) and (8), we can analyze the distribution of the fraction of admixture as a function of the time *g* and the parameters 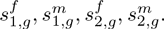 Under our model, *H*_1_*_,g,δ_* takes its values in *Q_g_* = {0, 1/2*^g^*, …, 1 − 1/2*^g^*, 1}. Therefore, using eqs. (7) and (8), and recalling that *H*_1_*_,g,f_* and *H*_1_*_,g,m_* are identically distributed, for a value *q* in the set *Q_g_*, we can compute the probability 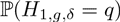 that a random individual from the hybrid population at generation *g* has admixture fraction *q*. For *g* = 1, 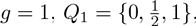 and

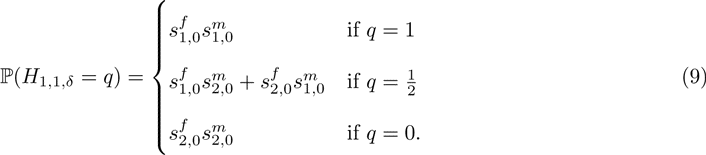

For all subsequent generations, *g ≥* 2, for *q* in *Q_g_*,

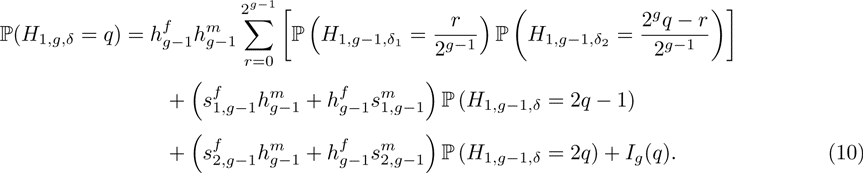

The function *I_g_* is defined for all values of *q* in *Q_g_*, an is equal to

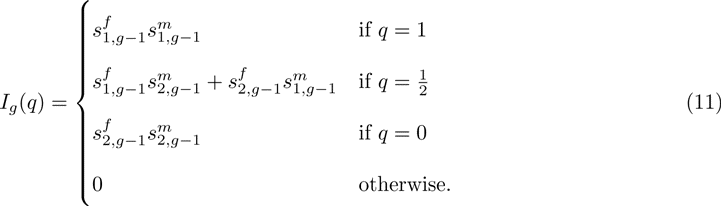

In eq. (10), we calculate the probability distribution of *H*_1_*_,g,δ_* by taking a sum over all possible parental pairings at the previous generation that would lead to an admixture fraction *q* at generation *g*. Only three values of *q* allow for a history without a single hybrid ancestor—*q* = 0, 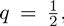 and *q* = 1—producing the terms in eq. (11). When there is no sex bias and 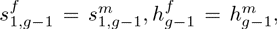 and 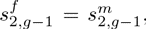 eqs. (9)–(11) reduce to the corresponding eqs. (3)–(5) from Verdu & Rosenberg (2011).

Eqs. (9)–(11) can be used to analyze the behavior of the distribution of *H*_1_*_,g−_*_1_*_,δ_* over time. In Figure 2,we consider constant admixture processes after the founding of the hybrid population 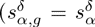 for each α ∈ {1,2}, δ ∈ {*f, m*}, and *g* ≥ 1) plotting 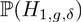 for the first six generations, as computed recursively using eq. (10). In Figure 2A and 2B, we consider a hybrid population founded with equal contributions from source populations *S*_1_ and *S*_2_, but with no further contributions after *g* = 1. In both of these cases, the distribution of the autosomal admixture fraction contracts around the mean of 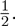 However, whereas Figure 2A has equal contributions from each sex in the founding generation, Figure 2B has a large initial sex bias. We see that the width of the distribution is smaller with the sex-biased contributions, despite equality of the total contributions *s*_1,0_ and *s*_2,0_.

**Figure 2:**
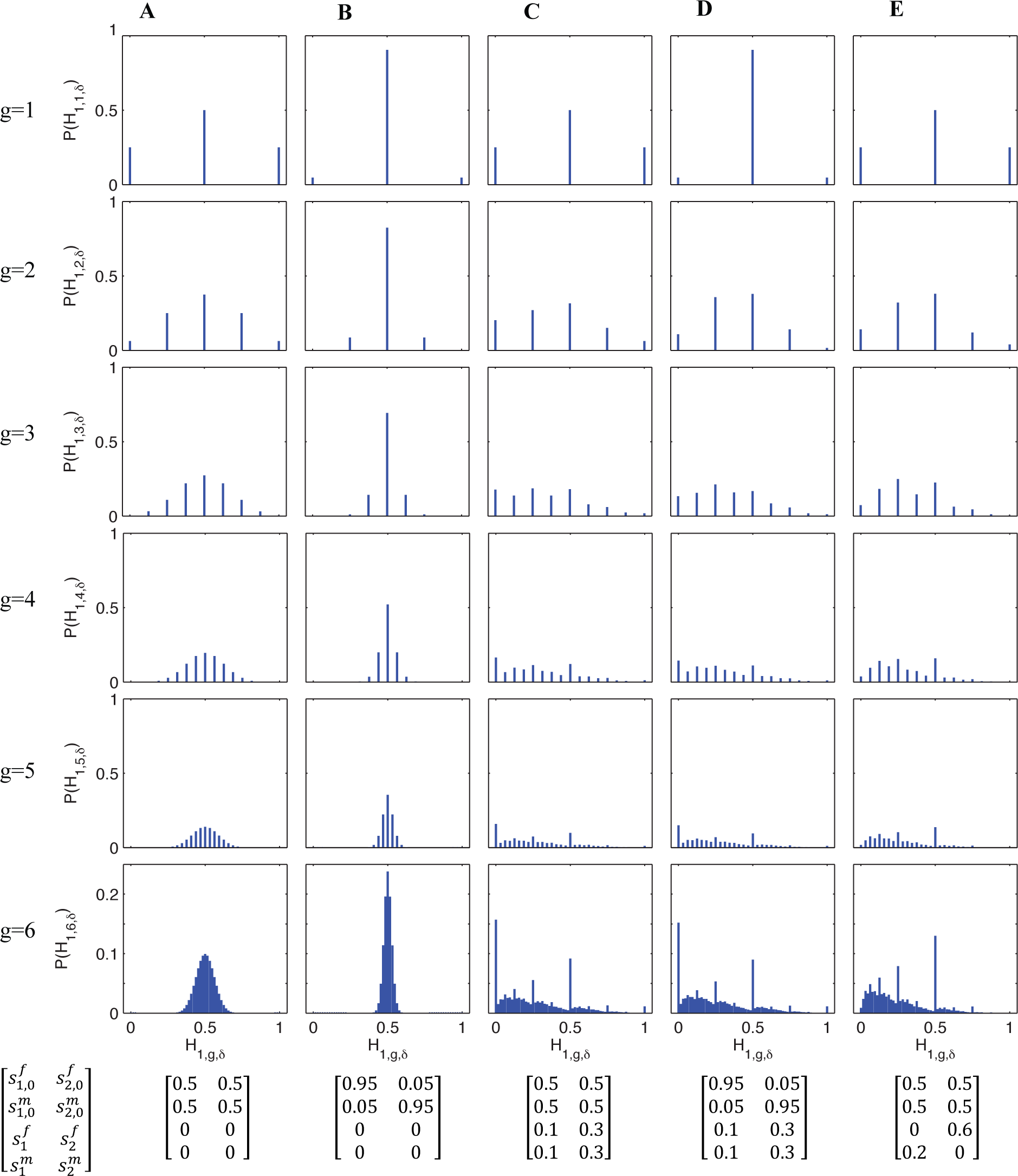
Probability distribution of the fraction of admixture from source population *S*_1_, 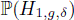 for a random individual from the hybrid population for the first six generations (eqs. (9)–(11)). Each column corresponds to a specified admixture scenario, with constant contributions from the source populations over time after founding 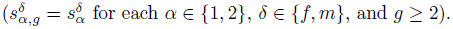

In Figures 2C-2E, we consider admixture scenarios in which the founding of the hybrid population is followed by constant contributions from the source populations over time, *s*_1_ = 0.1 and *s*_2_ = 0.3. Because the two source populations contribute after the founding, the distribution does not contract around the mean as in Figures 2A and 2B. Also, because the total contributions from *S*_1_ and *S*_2_ are unequal, the distribution of *H*_1_*_,g,δ_* is no longer symmetrical. Rather, because the contribution from *S*_2_ is greater, the distribution is shifted toward zero.

Figures 2C and 2D have the same continuing contributions for *g ≥* 2, with no sex bias in the founding generation for Figure 2C, and a large initial sex bias for Figure 2D. Despite different founding contributions, Figures 2C and 2D have similar distributions of *H*_1_*_,g,δ_* after a few generations. In Figure 2E, the hybrid population is founded without a sex bias and with equal contributions from the two source populations. The total contributions *s*_1_ and *s*_2_ are the same as in Figures 2C and 2D, but unlike in Figures 2C and 2D, the continuing contributions are sex-biased, with 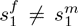 and 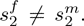 Even when *s*_1_ and *s*_2_ are held constant, the distribution of *H*_1_*_,g,δ_* depends on the *s^δ^_α_*. Notably, the probability of *H*_1,6,*δ*_ = 0 drops from 0.157 in Figure 2C to 0.000 in Figure 2E. Similarly, the probability of *H*_1,6,δ_ = 1 drops to zero in Figure 2E as well. With these reductions at the extremes, we see a rise in the probability of intermediate values for *H*_1*,g,δ*_.

## Expectation of the fraction of admixture

Using the law of total expectation, we write the expectation of the fraction of admixture from source population 1 for a random individual of sex *δ* in population *H* at generation *g* as a function of conditional expectations for all possible pairs of parents *L*,

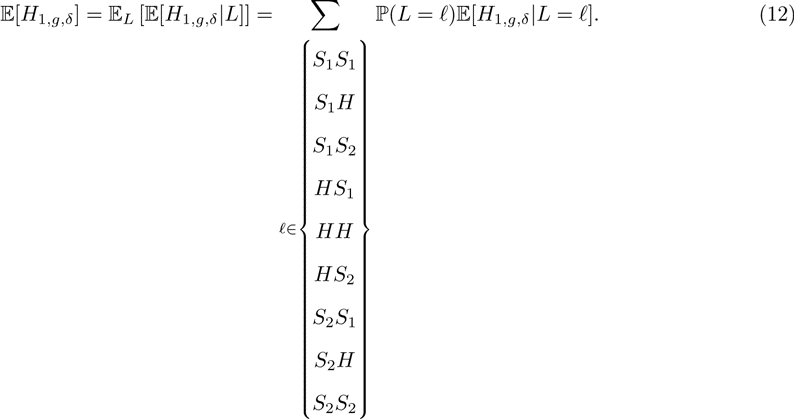

We can simplify this recursion relation. For *g* = 1,

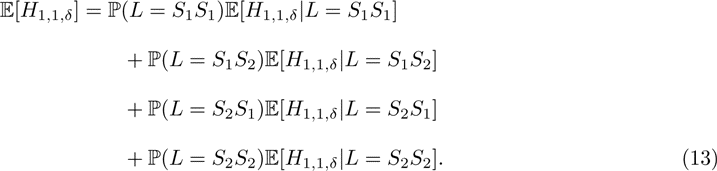

For all subsequent generations, *g ≥* 2, we have

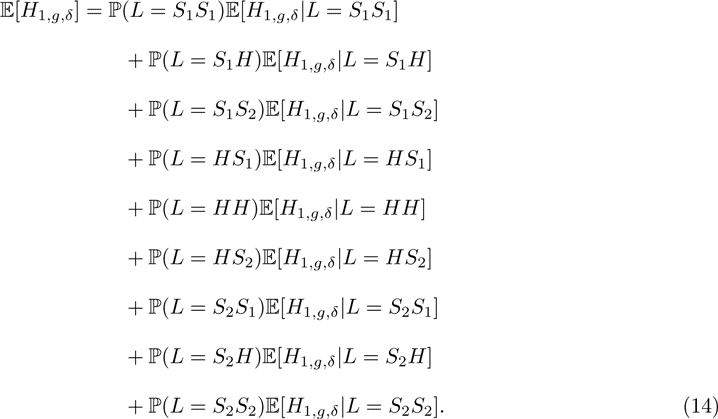

Using eqs. (7) and (8), for the first generation, *g* = 1, we have

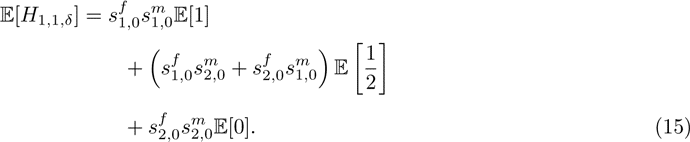

For all subsequent generations, *g ≥* 2, we have

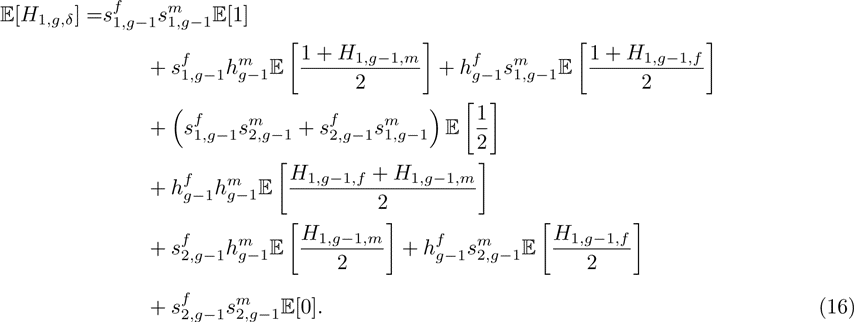

Recalling eqs. (1), (2), and (4), we can simplify the expectation of the fraction of admixture in a random individual of sex *δ* from the hybrid population. For *g* = 1, eq. (15) gives

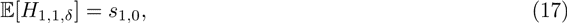

the same expression found by Verdu & Rosenberg (2011, eq. 10). For *g ≥* 2, by eq. (16),

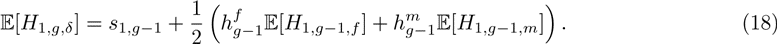

Because *H*_1_*_,g,f_* and *H*_1_*_,g,m_* are identically distributed, recalling eq. (2), we can simplify the expectation using 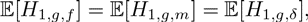 where *δ* is left as an unspecified sex (*f* or *m*). For *g ≥* 2, the expectation of the fraction of admixture from source population 1 is

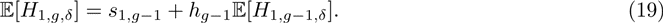

We see in eqs. (17) and (19) that the expectation of the fraction of admixture for a random individual of sex *δ* from the hybrid population at generation 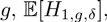 depends on the total contributions of the source populations (*S*_1_*, S*_2_*, H*) at each generation, *s*_1_*_,g−_*_1_ and *h_g−_*_1_, and not on the sex-specific parameters, 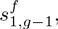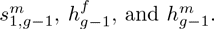 This recursion (eqs. (17) and (19)) is the same as in the non-sex-specific model of Verdu & Rosenberg (2011, eqs. 10 and 11).

## Higher moments of the fraction of admixture

We can write a general recursion for the higher moments of the fraction of admixture from population *S*_1_ in a randomly chosen individual of sex *δ* from the hybrid population. For *k ≥* 1, in the first generation, *g* = 1, we have

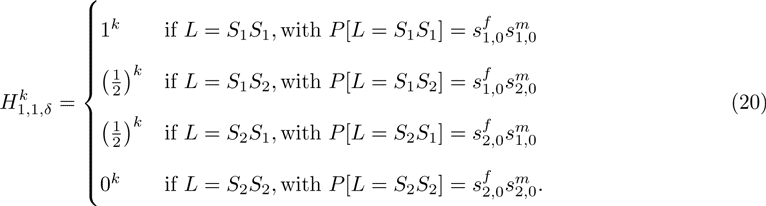

For all subsequent generations, *g ≥* 2,

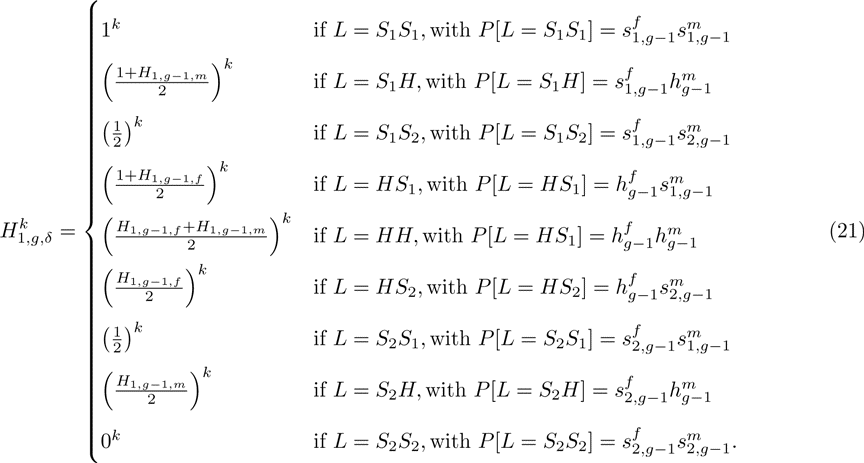

As in the case of *k* = 1, we use the law of total expectation to write a recursion for higher moments of the distribution of the fraction of admixture for all *k ≥* 1. Using the values for the recursion for the fraction of admixture, eqs. (7) and (8), for the first generation, *g* = 1, we have

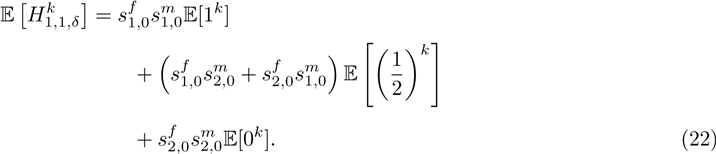

For *g ≥* 2, we have

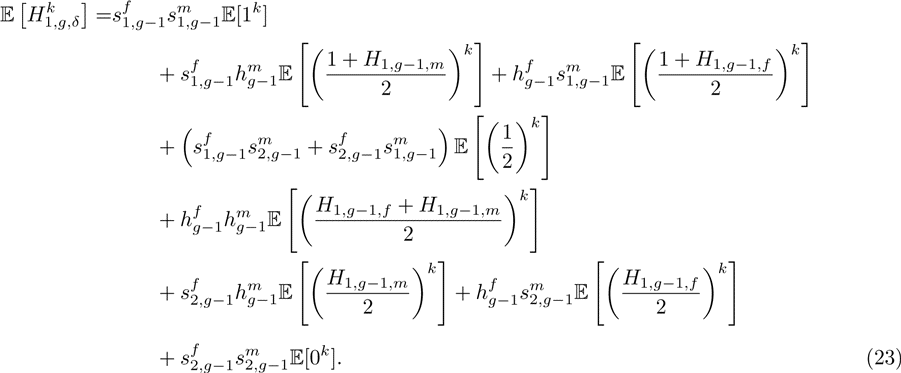

Recalling eq. (3) and noting that *h*_0_ = 0, we use the binomial theorem to simplify the recursion for the moments of *H*_1_*_,g,δ_*. For *g* = 1, we have

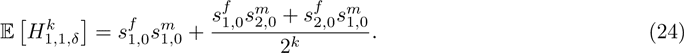

For *g ≥* 2, we have

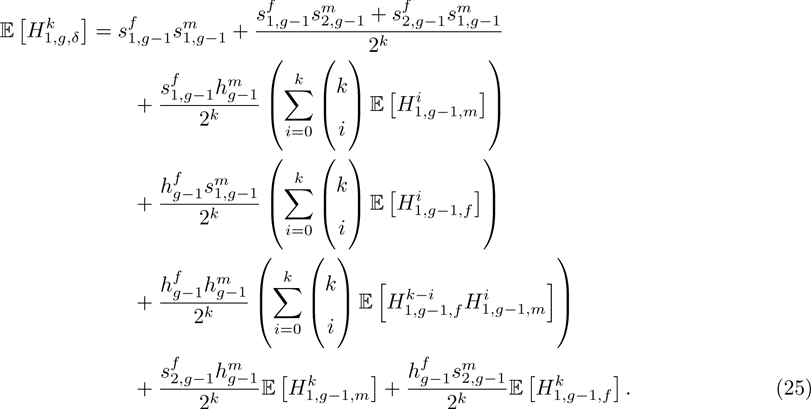

Because *H*_1_*_,g,f_* and *H*_1_*_,g,m_* have the same distribution, we can simplify the *k*th moment of the distribution of the fraction of admixture from *S*_1_, for *δ ∈* {*f, m*}, to give

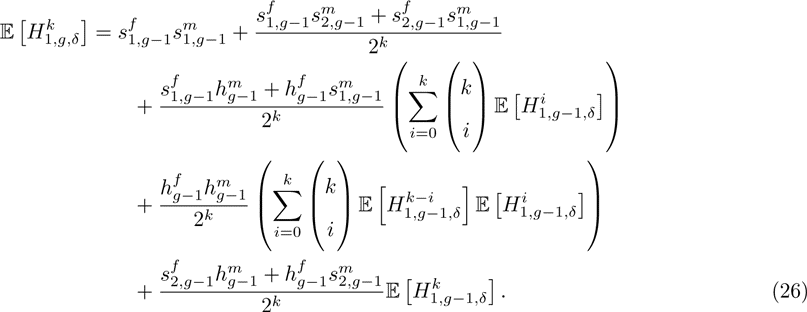

For *k* = 1, eqs. (24) and (26) should produce the expectation that we have already derived for *k* = 1. For *k* = 1, using eqs. (1), (2), and (4), eq. (24) gives

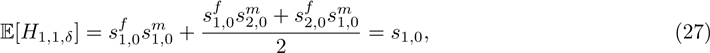

which matches eq. (17). For *g ≥* 2 and *k* = 1, eq. (26) gives

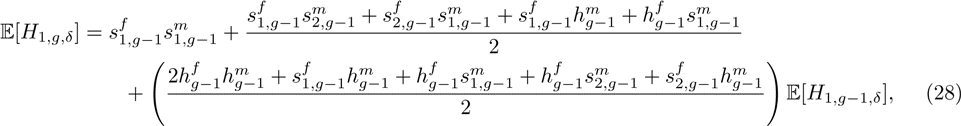

which simplifies to match eq. (19). Finally, with equal contributions in each population from females and males, so that 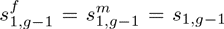 and 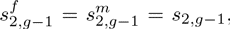 eqs. (24) and (26) reduce to eqs. 16 and 17 from Verdu & Rosenberg (2011).

## Variance of the fraction of admixture

When *k* = 2, eqs. (24) and (26) produce a recursion for the second moment of *H*_1_*_,g,δ_*. Recalling eqs. (1)–(6), for *g* = 1, we have

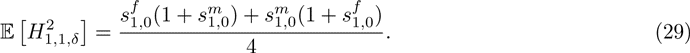

For *g ≥* 2, we have

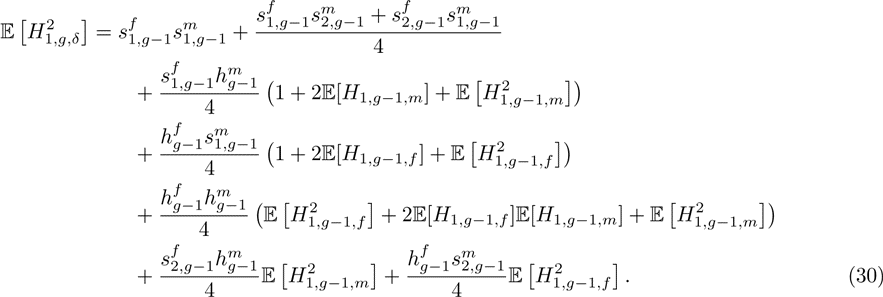

Recalling that *H*_1_*_,g,f_* and *H*_1_*_,g,m_* are identically distributed, eq. (30) simplifies to give

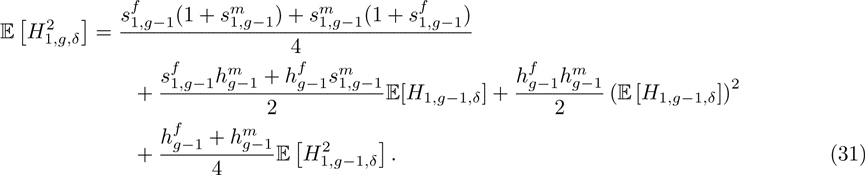

Using the definition of the variance 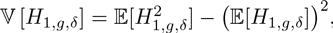 and eqs. (17), (19), (29) and (31), for the first generation, for the variance of the fraction of admixture, we have

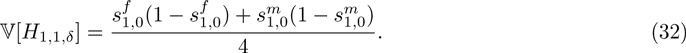

For all subsequent generations, *g ≥* 2, we have

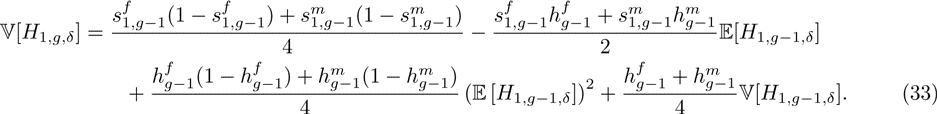

With no sex bias, so that 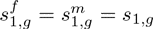 and 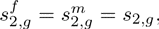 eqs. (32) and (33) are equivalent to eqs. 22 and 23 from Verdu & Rosenberg (2011).

The recursion for the variance of the fraction of admixture of a random individual of sex *δ* from the hybrid population is dependent on the variance from the previous generation, the expectation from the previous generation, and its square. By contrast with the expectation, the variance of the fraction of admixture depends on the sex-specific contributions from the source populations.

Eqs. (32) and (33) are invariant with respect to an exchange of all variables corresponding to males (superscript *m*) with those corresponding with females (superscript *f*). Thus, while the variance is affected by the sex-specific admixture contributions, it does not identify the direction of the bias. Despite the dependence of the variance of the autosomal fraction of admixture on sex-specific contributions, under the model, the symmetry demonstrates that autosomal DNA alone does not identify which sex contributes more to the hybrid population from a given source population. This result is reasonable given the non-sex-specific inheritance pattern of autosomal DNA.

## Special case: a single admixture event

Using the recursions in eqs. (17), (19), (32), and (33), we can study specific cases in which the contributions are specified. We first consider the case in which the source populations *S*_1_ and *S*_2_ do not contribute to the hybrid population after its founding: 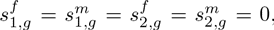 and 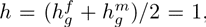 for all *g ≥* 1. As before, at the first generation, the hybrid population is not yet formed, and *h*_0_ = 0. Therefore, 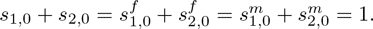

Under this scenario, we can derive the exact expectation and variance of the autosomal fraction of admixture of a random individual from the hybrid population. In the case of a single admixture event, the expectation of the admixture fraction is equal to the expectation at the first generation, because the further contributions are all zero. Using eq. (19), *s*_1,*g*−1_ = *s*_2_*_,g−_*_1_ = 0 for all *g ≥* 2. Therefore, from eq. (17), in the case of a single admixture event, for all *g ≥* 1,

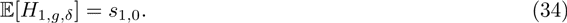

The expectation of the autosomal fraction of admixture from source population *S*_1_ is constant over time, and it depends on the total—not the sex-specific—contribution from the source population *S*_1_. As in the general case in eq. (19), for a single admixture event, a sex bias does not affect the expectation. Because the source populations provide no further contributions after the founding generation, unlike in the general case, the mean admixture fraction does not change with time.

Using eqs. (32) and (33), because 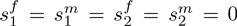 for all *g ≥* 2, the variance of the fraction of admixture follows a geometric sequence with ratio 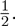 For all generations *g ≥* 1,

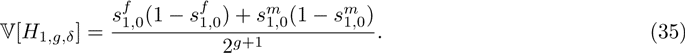

For 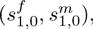 by eqs. (1) and (2), the variance matches eq. 25 of Verdu & Rosenberg (2011).

With a single admixture event, the variance decreases monotonically, and its limit is zero for all parameter values. Individuals from the hybrid population only mate within the population, decreasing the variance by a factor of two each generation. Thus, eq. (35) predicts that the distribution of the admixture fraction for a random individual in the hybrid population contracts around the mean, converging to a constant equal to the mean admixture from the first generation.

In eq. (35), considering all possible pairs 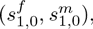 with each entry in [0, 1], the maximal 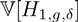 occurs at 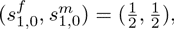 a scenario with equal cntributions from the two source populations, and no sex bias. At the maximum, the variance is 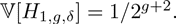 Four minima occur, at 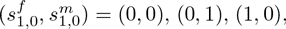 and (1, 1), cases in which all individuals in generation *g* = 1 have the same pair of source populations for their two parents, and in later generations, all individuals continue to have the same value of *H*_1_*_,g,δ_*. In these cases, 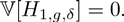

Figure 3 plots the variance in eq. (35) as a function of the sex-specific parameters 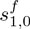 and 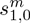 for three values of *g*. For *g* = 1, a maximum of 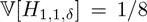 occurs at 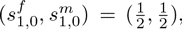 and a minimum, 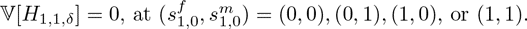 After one generation of mixing within the hybrid population, with no further contributions from the source populations, the maximum and minima occur at the same values of 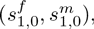 but the variance is halved (Fig. 3B). That is, for a given set of values 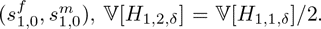 Similarly, for *g* = 8 in Figure 3C, 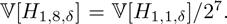 By *g* = 8, the hybrid population is quite homogeneous in admixture, and the variance of the admixture fraction has decreased to near zero for all sets of founding parameters. Therefore, the admixture fraction distribution is close to constant, with *H*_1,8_*_,δ_ ≈ s*_1,0_.

**Figure 3:**
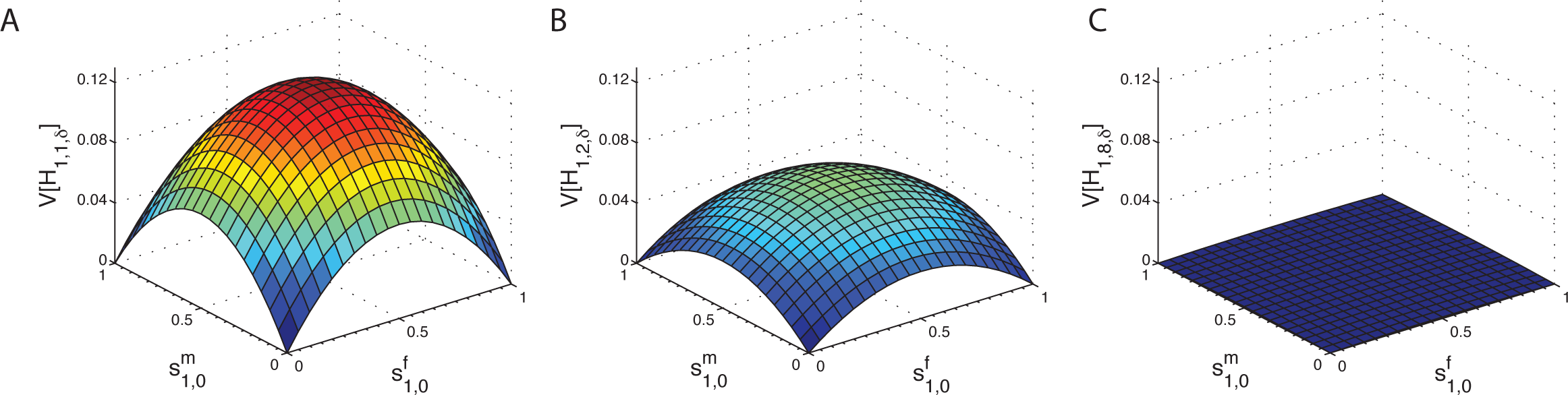
The variance of the fraction of admixture, *V* [*H*_1_*_,g,δ_*], as a function of female and male contributions from source population *S*_1_ in the first generation, 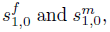 in the case of hybrid isolation. (A) *g* = 1. (B) *g* = 2. (C) *g* = 8. At each generation, the variance decreases toward zero by a factor of two. Considering all 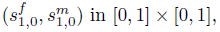 the maximal variance occurs when 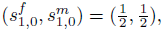 and the minimal variance occurs when 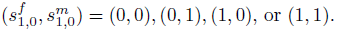 The variance is calculated using eq. (35).

We can analyze the dependence of the variance on the sex-specific parameters by considering constant total contributions *s*_1,0_ and allowing the sex-specific contributions to vary, constrained by eq. (1) so that 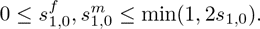 *≤* min (1, 2*s*_1,0_). Rewriting eq. (35) in terms of *s*_1,0_ and 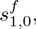

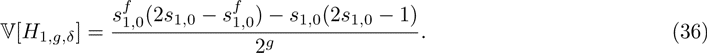

From this expression, it is possible to observe that given a constant *s*_1,0_ in [0, 1], the maximal variance is produced when 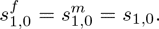 The minimal variance occurs when 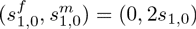 or (2*s*_1,0_, 0) for 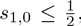 or (1, 2s_1,0_ − 1) or (2s_1,0_ −1,1) for 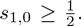 This minimum only takes the value 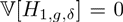 when *s*_1,0_ equals 0, 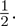 or 1.

For the specific case of 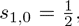 the total contribution for which the maximal variance occurs in Figure 3, we illustrate the variance at several locations in the allowed range for 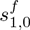 and 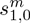 (Fig. 4). Four scenarios are plotted with the same total founding contribution from source population 1, 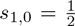 but with different levels of sex bias. As the female and male contributions become increasingly different, the initial variance decreases. The largest variance for 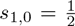 occurs at 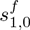 = 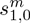 = *s*_1,0_, with no sex bias. The minimum occurs when males all come from one source population and females all from the other. In this extreme sex-biased case, the variance is zero constantly over time, as each individual has a male parent from one population and a female parent from the other, and an admixture fraction of 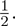

**Figure 4:**
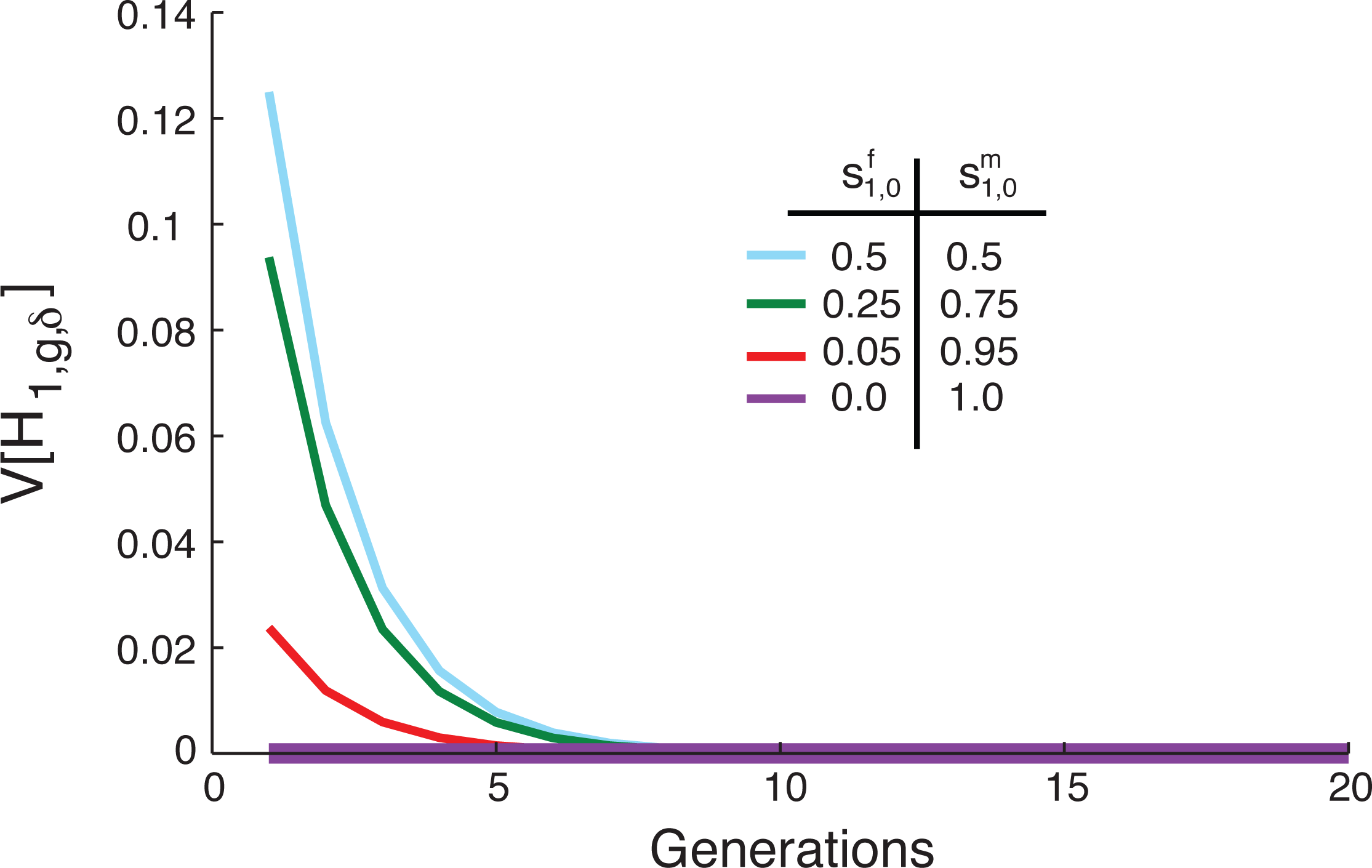
The variance of the fraction of admixture, *V* [*H*_1_*_,g,δ_*], when contributions from the source populations occur only in the founding generation and the total contribution from source population 1 is held constant at 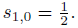 The limit of the variance of the fraction of admixture over time is zero for any choice of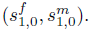 The magnitude of the variance, calculated from eq. (35), is inversely related to the level of sex bias. For all four scenarios, *s*_1_ = *s*_2_ = 0 and 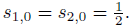

## Special case: constant nonzero contributions

Next, we consider the case in which an initial admixture event founds the hybrid population, and is then followed by constant nonzero contributions from the source populations. After the founding, for each *g ≥* 1, all admixture parameters are constant in time: 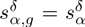 for each *α* ∈ {1, 2} and *δ* ∈ {f, m}, and 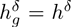 for each *δ*. Thus, we have parameter values for the founding, and constant continuing admixture parameters 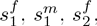 and 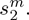 Each parameter takes its value in [0, 1], as do *s*_1_ and *s*_2_. By contrast, *h* takes its value in (0, 1). The case of *h* = 1 is a single admixture event, analyzed above. The *h* = 0 case is trivial because the hybrid population is re-founded at each generation, and the distribution of the admixture fraction thus depends only on the contribution in the previous generation. Therefore, we require *s*_1_+*s*_2_ ≠ 0 and *s*_1_+*s*_2_ ≠ 1.Individually, however, *h^f^* and *h^m^* can each vary in [0, 1], as long as they are not both zero or one.

The recursion for the expectation of the autosomal fraction of admixture, eqs. (17) and (19), is equivalent to that derived by Verdu & Rosenberg (2011). Therefore, the closed form of the expectation is equivalent as well. From Verdu & Rosenberg (2011) eq. 30, we have

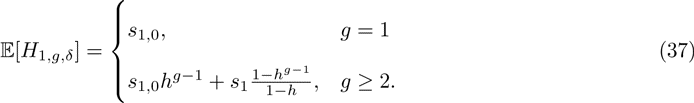

We can use the same method as Verdu & Rosenberg (2011) to simplify the second moment. Under the special case of constant contributions across generations, for *g* = 1, eq. (29) gives

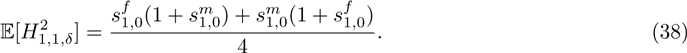

For g ≥ 2, eq. (31) gives

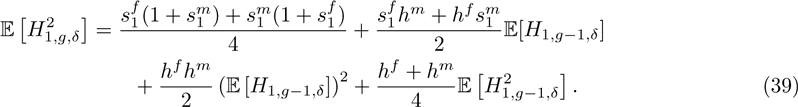

Because this equation is a non-homogenous first-order recurrence with the form

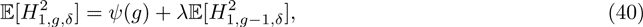

we can use Theorem 3.1.2 of Cull et al. (2005) to solve for a unique solution for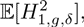 as in Verdu & Rosenberg (2011). For the initial condition, we have

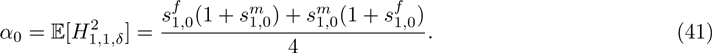

We define *λ* = *h^f^* + *h^m^ /*4 = *h/*2, and for all *g ≥* 2, we have

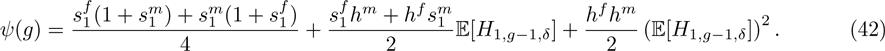

Using the expected admixture fraction from eq. (37), we can simplify eq. (42). For all *g ≥* 2,

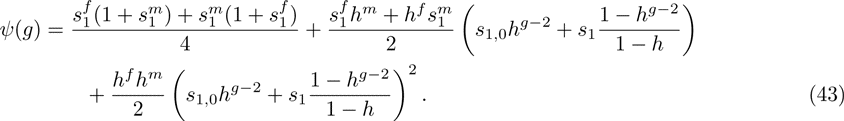

Therefore, using Theorem 3.1.2 of Cull et al. (2005), we have a unique solution for 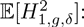

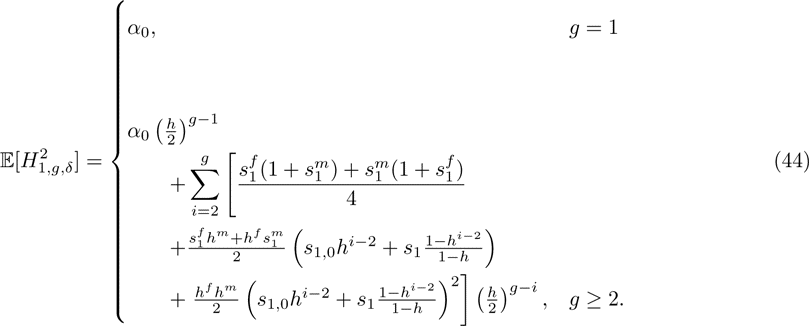

Eq. (44) can be simplified by separating the sum and summing the resulting geometric series:

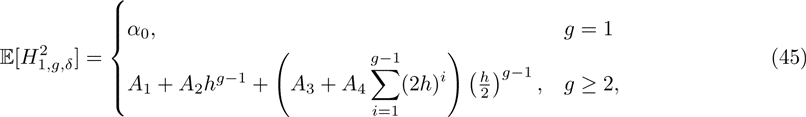

where *α*_0_ is defined in eq. (41), and

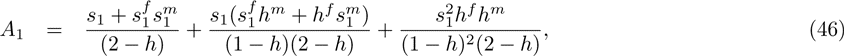

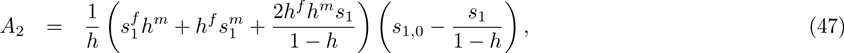

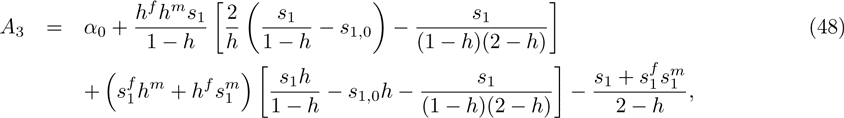

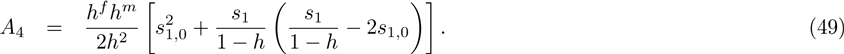

When 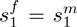 and 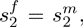 *A*_1_, *A*_2_, *A*_3_, and *A*_4_ are equal to the corresponding quantities in eqs. 39–42 of Verdu & Rosenberg (2011). Therefore, without sex bias, the closed form of the second moment of the admixture fraction, eq. 45, is equal to eq. 38 in Verdu & Rosenberg (2011).

Using the relation 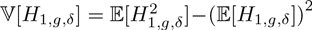 and eqs. (37) and (45), for the variance of the autosomal fraction of admixture, we have

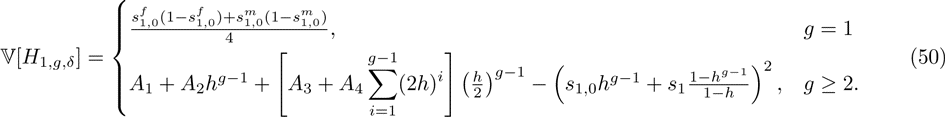

For 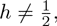 we have

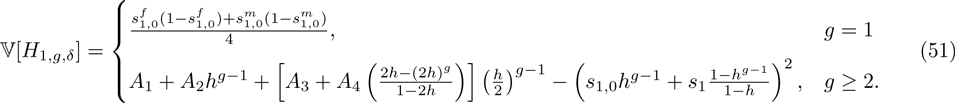

For 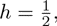 eq. (50) gives

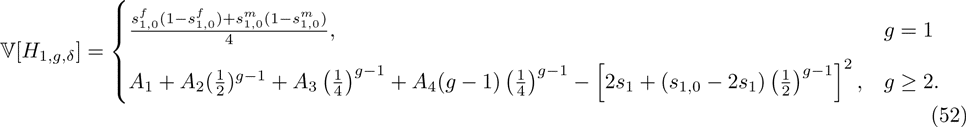

Eqs. (50)–(52) simplify to eqs. 43–45 of Verdu & Rosenberg (2011) when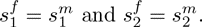

### Limiting variance of admixture over time

Figure 5 illustrates the variance of the autosomal fraction of admixture as a function of *g* when the contributions from the source populations are constant over time, computed using eq. (50). The figure shows that if the continuing contributions are held constant, then the long-term limiting variance does not depend on the founding parameters. Unlike in the hybrid isolation case, under the scenario of constant, nonzero contributions from the source populations over time, *h* ≠ 0 and *h* ≠ 1, a nonzero limit is reached. Applying eq. (50), we have

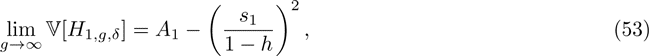

which does not depend on the founding parameters. The limit matches that of Verdu & Rosenberg (2011, eq. 46) in the absence of sex bias 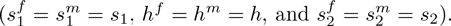

**Figure 5:**
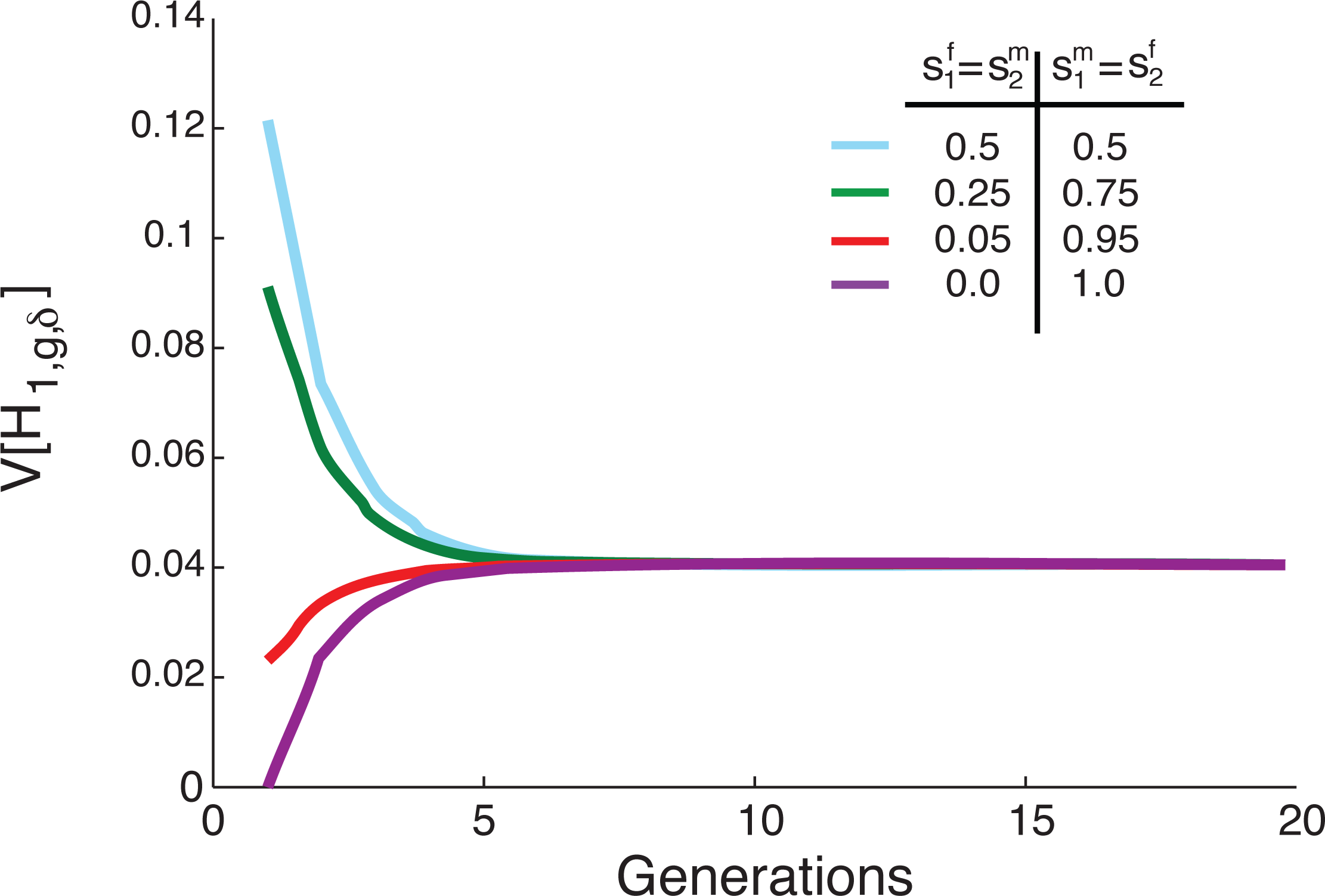
The variance of the fraction of admixture over time for constant, nonzero contributions from the source populations, with different levels of sex bias in the founding of the hybrid population, and constant, equal, and nonzero subsequent contributions from the source populations and sex for *g ≥* 1. In all cases, 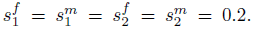 The variance, calculated with eq. (50), reaches a nonzero limit 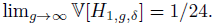

### The maxima and minima of the limiting variance

Using eqs. (1), (2), (4) and (46), the limit in eq. (53) can be equivalently written in terms of the two female sex-specific contributions, 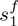 and 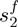 and the total contributions from the two source populations, *s*_1_ and *s*_2_. Considering admixture scenarios with constant *s*_1_*, s*_2_, with *s*_1_ + *s*_2_ *∈* (0, 1], but allowing 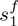 and 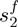 to range over the closed unit interval, the limiting variance depends on two independent parameters, 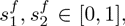 subject to the constraint in eq. (1):

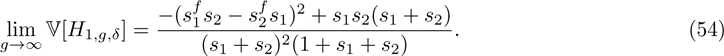

Treating *s*_1_ and *s*_2_ as constants in [0,1], the critical points of eq. (54) are the same as those of

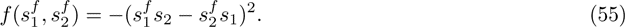

First we consider the maximum. Because 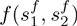 is always negative or zero, the maximal variance given *s*_1_ and *s*_2_ occurs when 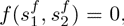 which occurs on the line 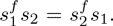 Equivalently, recalling eq. (1), this line can be written as

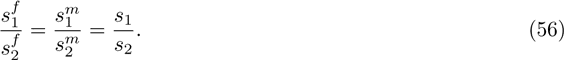

Eq. (56) has many solutions for 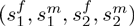 given *s*_1_ and *s*_2_. One solution is 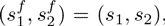 which by eq. (1) is equivalent to 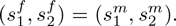 Therefore, the limiting variance of the admixture fraction is maximized when there is no sex bias. Figure 6 plots two examples of the variance for constant *s*_1_ and *s*_2_, but increasingly different sex-specific contributions from the source populations. In both panels, the admixture history with no sex bias produces the greatest limit.

**Figure 6:**
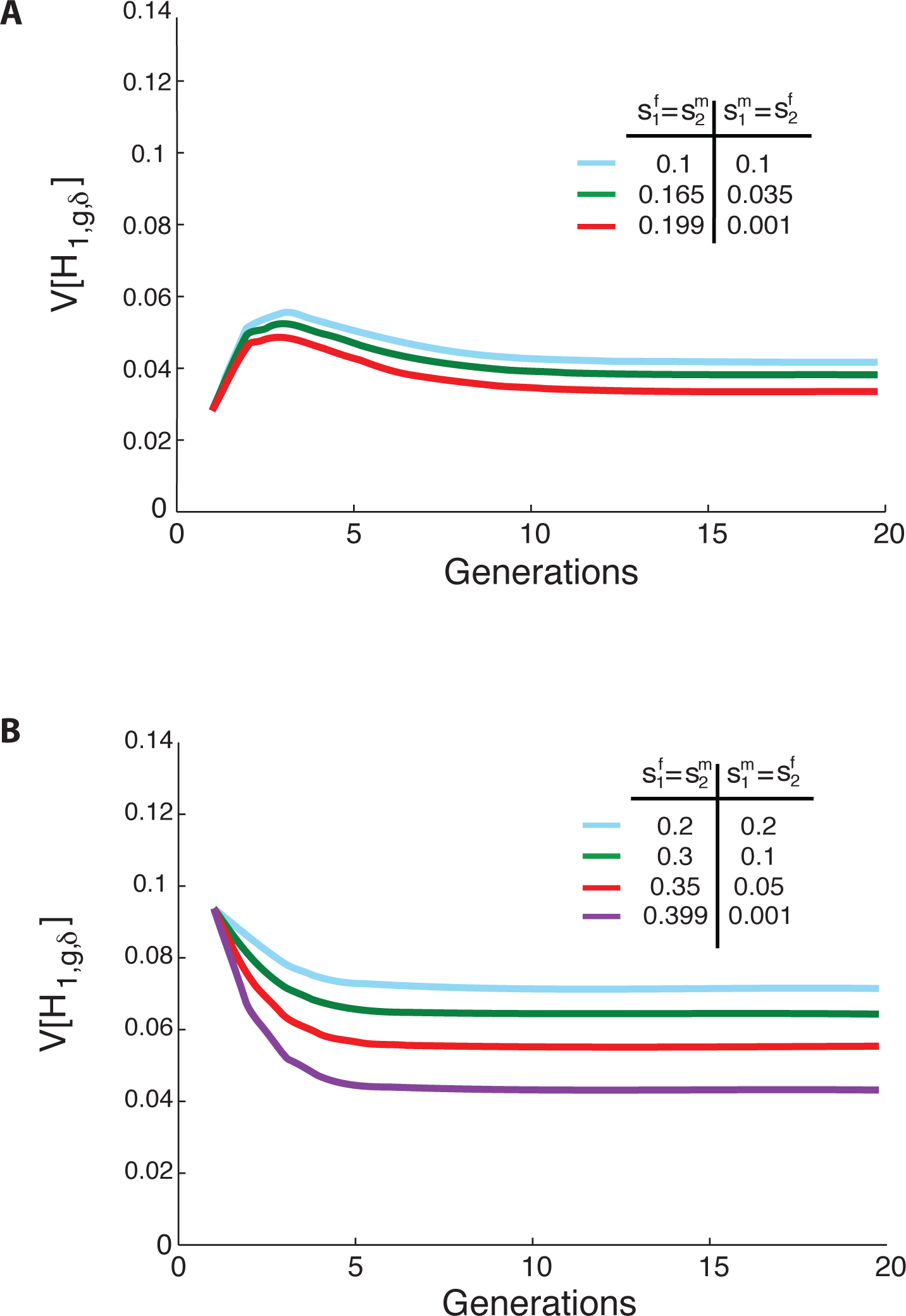
The variance of the fraction of admixture over time for constant, nonzero contributions from the source populations, with different levels of sex bias, but the same total contribution from the two source populations. (A) 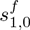 = 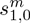 = 0.9, and *s_1_* = *s_2_* = 0.1, (B) 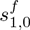 = 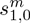 = 0.75, and *s_1_* = *s_2_* = 0.2 The variance reaches a nonzero limit when 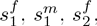 and 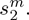 are nonzero and constant over time. The variance is calculated using eq. (50).

For fixed *s*_1_ and *s*_2_, however, the case without sex bias is not the only maximum of the limiting variance. Figure 7 plots the variance over time for four different admixture histories, each with the same total contributions *s*_1_ and *s*_2_, but quite different sex-specific contributions 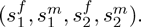 Each of the four scenarios plotted reaches the same limit because each provides a solution to eq. (56). Because 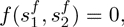 eq. (54) depends only on the total contributions *s*_1_ and *s*_2_. For constant *s*_1_ and *s*_2_, any admixture history whose contributions solve 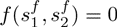 has limiting variance

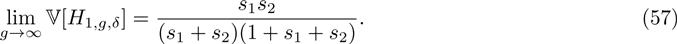

This maximal limiting variance depends on the total contributions from the source populations, but not on the sex-specific contributions; it is equivalent to eq. 47 of Verdu & Rosenberg (2011).

**Figure 7:**
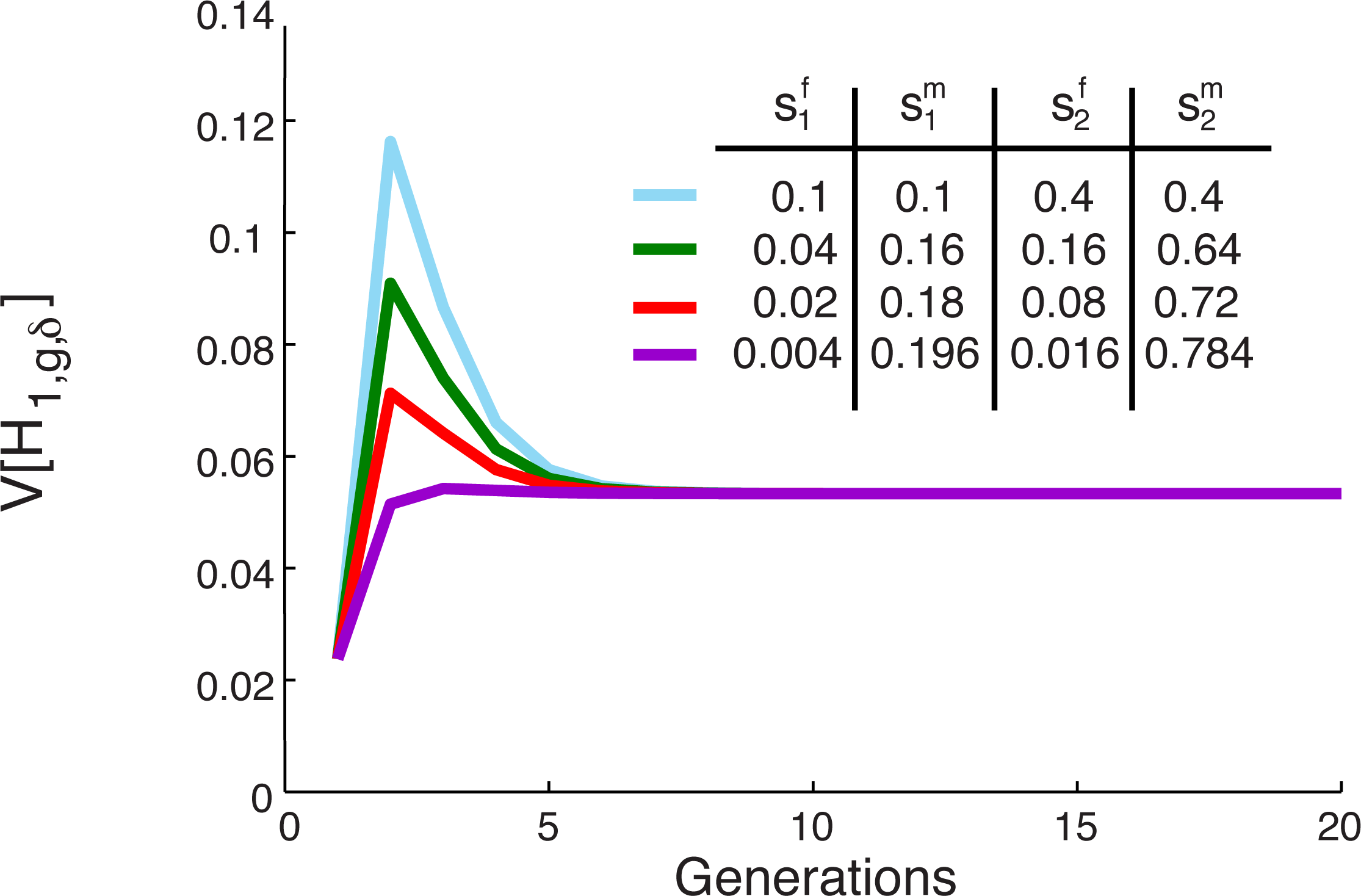
The variance of the fraction of admixture over time for constant, nonzero contributions from the source populations, but multiple different ratios of female to male contributions. When the sex-specific parameters satisfy the equation 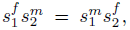 multiple different demographic scenarios have the same limiting variance of the admixture fraction. In all cases,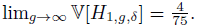 The variance is calculated using eq. (33). For all scenarios 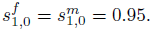

Thus far, we have considered the maximal limiting variance as a function of the sex-specific parameters given constant total contributions *s*_1_ and *s*_2_. We can also identify the values of *s*_1_ and *s*_2_ that maximize the limiting variance, considering all *s*_1_*, s*_2_ *∈* [0, 1]. For each choice of *s*_1_ and *s*_2_, the maximal variance over values of 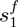 and 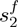 is given by eq. (57). We can therefore find the *s*_1_ and *s*_2_ that maximize eq. (57). As demonstrated by Verdu & Rosenberg (2011), given *s*_1_ + *s*_2_, the maximal limiting variance occurs when *s*_1_ = *s*_2_. Over the range of possible choices for *s*_1_ + *s*_2_ *∈* (0, 1), the maximum occurs when 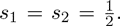 Unlike in Verdu & Rosenberg (2011), however, this maximum requires the sex-specific contributions to solve 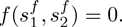

Interestingly, one of the *minima* of the limiting variance occurs when 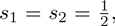 but with 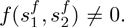 Specifically, when 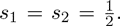 but all males come from one source population and all females from the other, 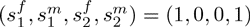, the limiting variance in eq. (54) is zero. In this case, *L* = *S*_1_*S*_2_ or *L* = *S*_2_*S*_1_ for every individual in the hybrid population. By eq. (8), the hybrid population is founded anew at each generation, with each individual having admixture fraction 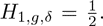 Therefore, the population has zero variance.

More generally, given *s*_1_ and *s*_2_, the minimal limiting variance occurs when 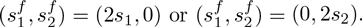 Given *s*_1_ and *s*_2_, the limiting variance is minimized with respect to 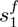 and 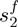 when 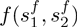 is smallest (eq. (54)). Because *f* is the negative of the square of a difference of products, it is greatest when one term is zero and the other is at its maximum, as at 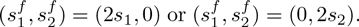 These points represent the maximal sex bias for fixed *s*_1_, *s*_2_.

If we allow *s*_1_ and *s*_2_ to vary, because a variance is bounded below by zero, any set of parameters that produces zero variance is a minimum. In eq. (54) if either *s*_1_ = 0 or *s*_2_ = 0, then the limiting variance of the admixture fraction is zero. When only one population contributes after the founding, in the limit, all ancestry in the hybrid population traces to that population.

### Properties of the limiting variance

The limiting variance of the fraction of admixture over time in eq. (53) is a function of the sex-specific contributions from the hybrid population, *h^f^* and *h^m^*, and source population 1, 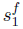 and 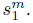 Recalling eq. (4), the limiting variance is equivalently written as a function of the sex-specific contributions from source population 2, *s*_2_^*f*^ and *s*_2_^*m*^, and either source population 1 (eq. (54)), or the hybrid population. It can be viewed as a function of all six sex-specific parameters 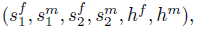 four of which can be selected while assigning the other two by the constraint from eq. (4).

We can therefore analyze the behavior of the limiting variance as a function of two of the sex-specific parameters by specifying two other parameters, and allowing the final two parameters, one female and one male, to vary according to eq. (4). Of the four parameters we consider, using the constraint from eq. (4) separately in males and females, two must be male and two must be female. Because the variance is invariant with respect to exchanging the source populations or the sexes, the six-dimensional parameter space generates a number of symmetries. Figures 8–12 examine the five possible, non-redundant ways of choosing two populations and the corresponding male and female parameters from those populations, and holding two corresponding parameters fixed (either from the same sex in the two populations, or for males and females from one population) while allowing the other two to vary. Figure 13 then highlights an informative case that considers the limiting variance as a function of a male and a female parameter from different populations.

**Figure 8:**
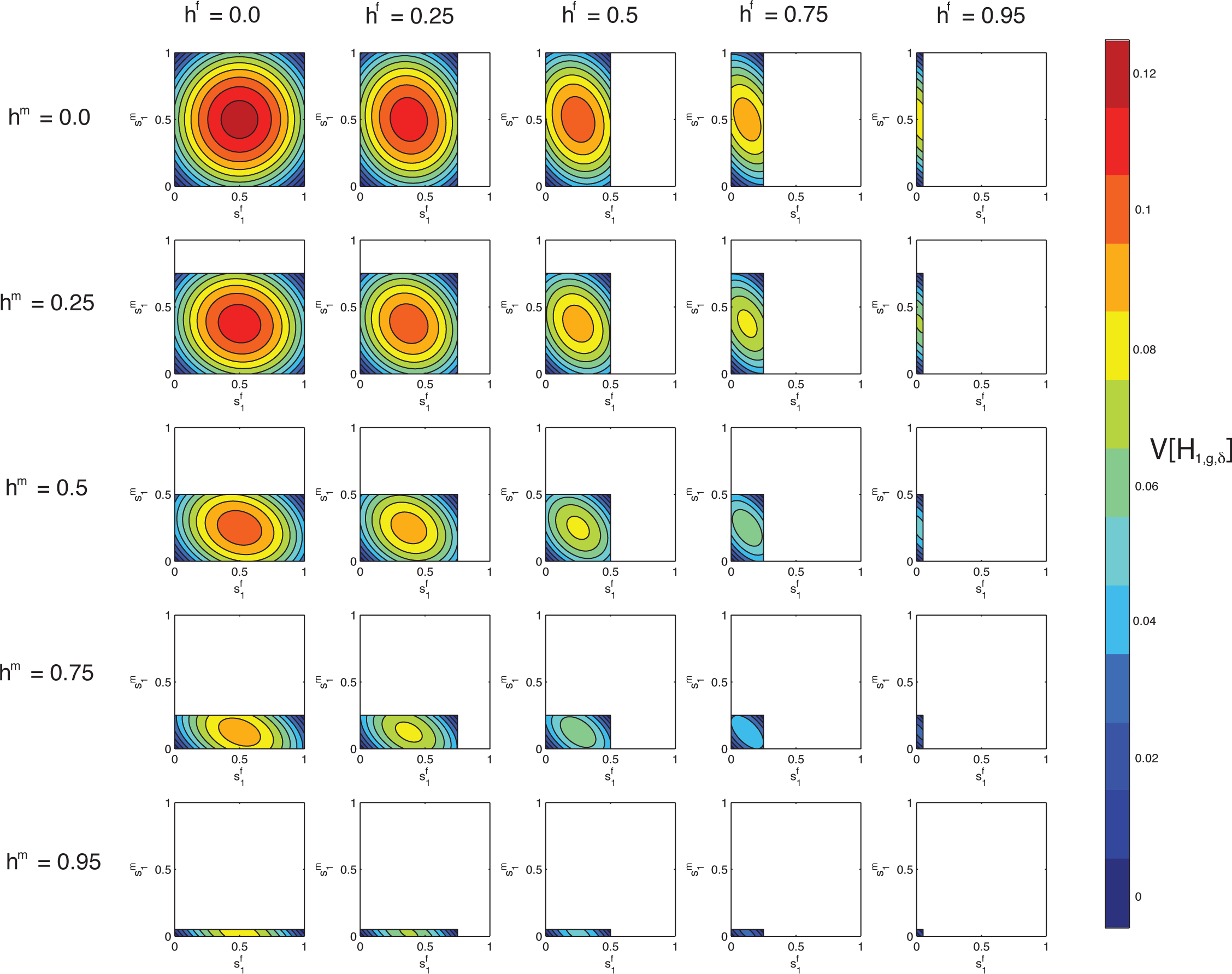
Contour plots of the limit of the variance of the fraction of admixture over time as a function of *s*_1_^*f*^ on the x-axis and *s*_1_^*m*^ on the y-axis for specified values of *h*^*f*^ by column and *h*^*m*^ by row. The domains of *s*_1_^*f*^ and *s*_1_^*m*^ are [0,1−*h^f^*] and [0,1−*h^m^*] respectively.

Each figure shows multiple contour plots of the limiting variance as a function of two sex-specific parameters, for fixed values of two other parameters. Three cases plot the limiting variance as a function of the female and male parameters from a given population, with the female and male contributions of another population specified. In two other cases, parameters for a single sex from two populations are plotted, specifying the contributions from the other sex for those populations.

By considering these parameter combinations, we can examine the dependence of the variance on sex-specific parameters and parameter interactions, as well as potential bounds on both the parameters and the variance. We highlight a number of symmetries in the limiting variance. The plots also illustrate the maxima and minima found in the previous section.

### Properties of the limiting variance in terms of 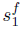 and 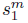

In each panel in Figure 8, we consider the variance of the fraction of admixture as a function of 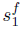, the female contribution from *S_1_*, on the x-axis, and 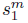, the male contribution from *S*_1_, on the y-axis, computed using eq. (53). We plot the variance for fixed *h^f^*, the female contribution from *H*, and *h^m^*, the male contribution from *H*. The domain for 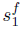 and 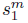 is constrained by eq. (4), with 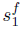 taking values in [0, 1 *− h^f^*], and 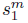 taking values in [0, 1 *− h^m^*].

The upper left plot in Figure 8 shows the variance as a function of 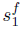 and 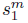 with *h^f^* = *h^m^* = *h* = 0. In this setting, the hybrid population is founded anew by the source populations each generation, and 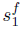 and 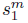 both take values from the full domain [0, 1]. For *h^f^* = *h^m^* = *h* = 0, the maximal limiting variance is lim_*g* → ∞_ 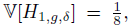 occurring when 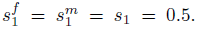 At this maximum, given eq. (4) and *h^f^* = *h^m^* = 0, we have 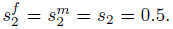 Therefore, as in eq. (54), the maximal limiting variance occurs when female and male contributions from the source populations are equal, and the total contributions from the source populations are equal.

The minima of lim_*g* → ∞_ 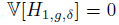 occur at the four corners of the plot. At the origin, when 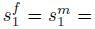 *s*_1_ = 0, the limiting variance is zero because only *S*_2_ contributes to the hybrid population. Individuals in the hybrid population all have parents *L* = *S*_2_*S*_2_, and an admixture fraction of zero (eq. (8)). By exchanging *S*_1_ for *S*_2_, the case of 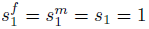 is similar.

Additional minima occur at 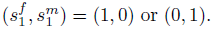 Here, all males come from one source population and all females from the other. Therefore, all individuals at the next generation of the hybrid population have parents *L* = *S*_1_*S*_2_ or *L* = *S*_2_*S*_1_, and admixture fraction 1/2 (eq. (8)).

For *h^f^* = *h^m^* = *h* = 0, the limiting variance is symmetrical over the line 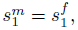 as a result of the symmetry between males and female in the variance (eq. (32)). Because the hybrid population provides no contribution and the variance of the fraction of admixture is symmetric with respect to source population, the variance is also symmetric over the lines 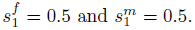

The columns of Figure 8 consider increasing, fixed values for *h^f^*, and the rows consider increasing, fixed values for *h^m^*, both from {0, 0.25, 0.5, 0.75, 0.95}. All panels maintain the general shape of the limiting variance as a function of 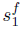 and 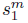 seen for *h^f^* = *h^m^* = 0. However, as the domain for 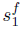 and 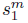 shrinks with increasing *h^f^* and *h^m^*, the location of the maximal variance changes across panels. In all cases, the maximum of the limiting variance occurs when 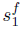 and 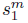 each lie at the midpoints of their respective domains, 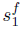 = (1 −*h^f^*)/2 and 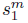= (1 − *h^m^*)/2). The magnitude of the limiting variance at each maximum decreases as its location moves away from 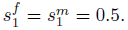

For each panel, the minimum lim_*g* → ∞_ 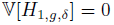 occurs when 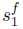 and 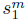 are either both zero or they lie at the maxima of their respective domains. In these cases, only one source population contributes to the hybrid population, and therefore, all individuals in the hybrid population have an admixture fraction from *S*_1_ of either 0, when 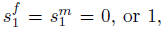 when 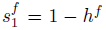 and *s*_1_^*m*^ = 1−*h^m^* ((8)). The limiting variance is no longer zero at the two corners of each contour plot where only one of {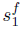, 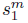} is at the maximum of its domain; these corners, however, are minima of the variance conditional on the values of *s*_1_ and *s*_2_. In these cases, males all come from one source population and females from the other, producing a minimum of eq. (54) conditional on fixed *s*_1_ and *s*_2_.

As in the case of *h^f^* = *h^m^* = 0, each plot is symmetrical in reflecting over both the midpoint of the x-axis, 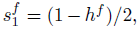 and that of the y-axis, 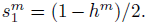 The limiting variance is symmetrical with respect to source population (eq. (54)), and this pair of reflections corresponds to an exchange of source populations. For *h^f^* = *h^m^* = 0, the line 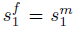 generates circular contours, but as the contributions from the hybrid population increase, the contours become elliptical.

In Figure 8, plots on the diagonal have equal contributions from males and females in the hybrid population, *h^f^* = *h^m^* = *h*. For *h^f^* ≠ *h^m^,* plots above the diagonal are equivalent to those below the diagonal with an exchange of female for male contributions, for both *s*_1_ and *h*. For example, the plot with *h^f^* = 0.25 and *h^m^* = 0.5 is equivalent to the plot with *h^f^* = 0.5 and *h^m^* = 0.25 if the axes are also switched so that 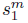 appears along the x-axis and 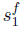 is on the y-axis.

Figure 9 plots the limiting variance as a function of *s*_1_^*f*^ on the x-axis, and *s*_1_^*m*^ on the y-axis, as in Figure 8, but we now fix values of 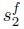 by column and 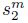 by row using eqs. (54) and (1). The maxima and minima occur at the same parameter values found in Figure 8, but they appear in different locations on the plots. For example, in Figure 9, the global maximum across panels occurs in the plot with 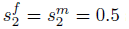 specified, and 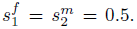 By eq. (4), this location implies *h^f^* = *h^m^* = 0, the plot that contains the maximal variance in Figure 8. In the upper left plot in Figure 9, the limiting variance is a constant zero for all *s_1_^f^* and *s_1_^m^* because *s*_2_ = 0 (eq. (54)).

**Figure 9:**
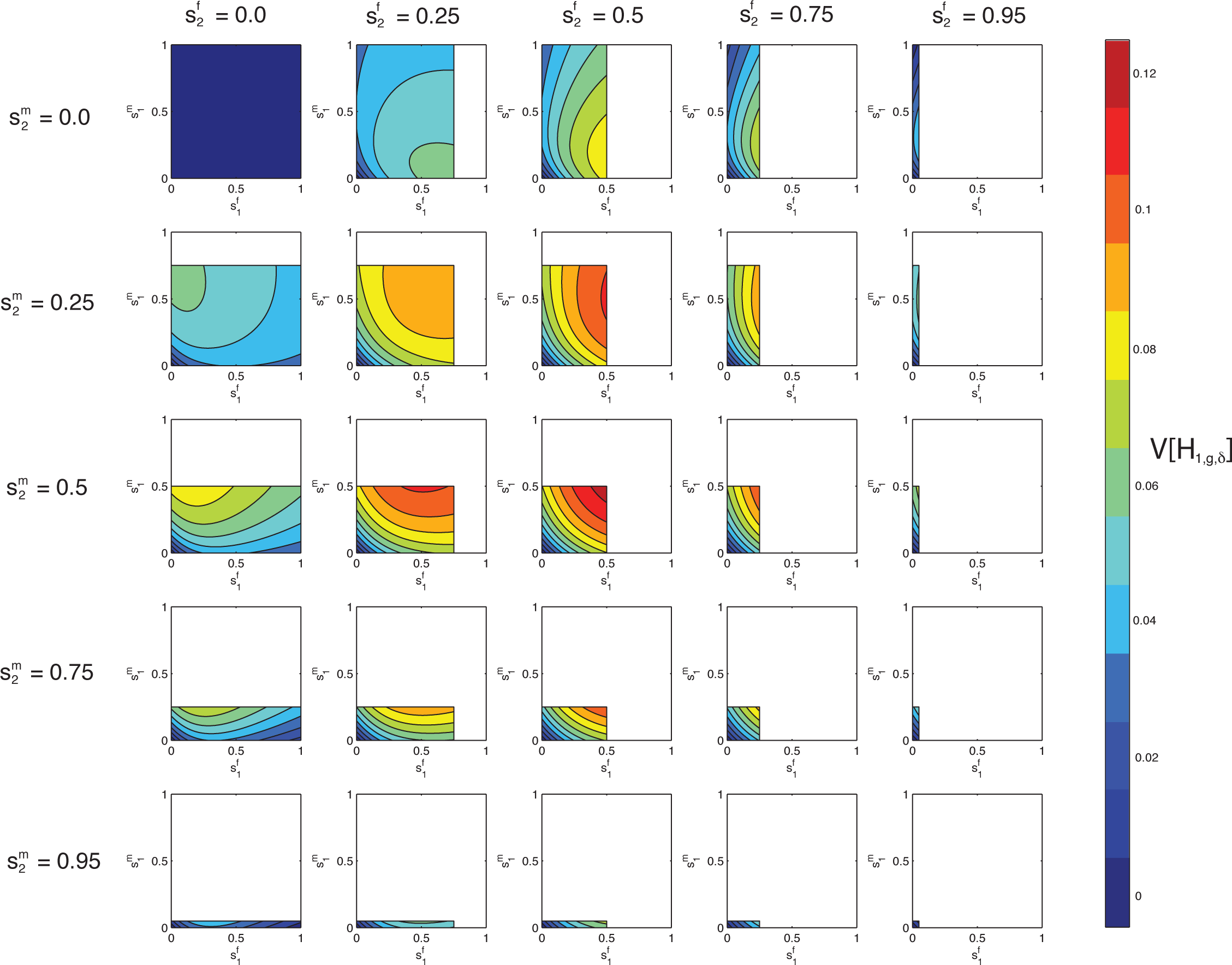
Contour plots of the limit of the variance of the fraction of admixture over time as a function of *s*_1_^*f*^ on the x-axis and *s*_1_^*m*^ on the y-axis for specified values of *s*_2_^*f*^ by column and *s*_2_^*m*^ by row. The domains of *s*_1_^*f*^ and *s*_1_^*m*^ are 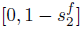 and 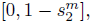 respectively.

Whereas all panels in Figure 8 are symmetric in reflecting over the midpoints of both domains, in Figure 9, only the plots with 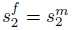 are symmetric over the line 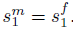 However, the symmetry corresponding to transposing males and females is visible in that a plot above the diagonal and its corresponding plot below the diagonal are equivalent if the axes for *s_1_^f^* and *s_1_^m^* are switched.

### Properties of the limiting variance in terms of *h^f^* and *h^m^*

Similarly to Figure 8, Figure 10 considers the limit of the variance of the fraction of admixture over time as a function of the four variables 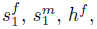 and *h^m^* using eq. (53). In Figure 10, each plot shows *h^f^* on the x-axis and *h^m^* on the y-axis, for specified values of *s_1_^f^* and *s_1_^m^* with the domains of *h^f^* and *h^m^* constrained by eq. (4). As in Figure 8, the plots along the diagonal are the cases with 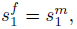 and there is a symmetry over this line of plots in that if the values of 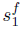 and 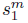 are switched, then the plots will be equivalent with a transposition of the axes.

**Figure 10:**
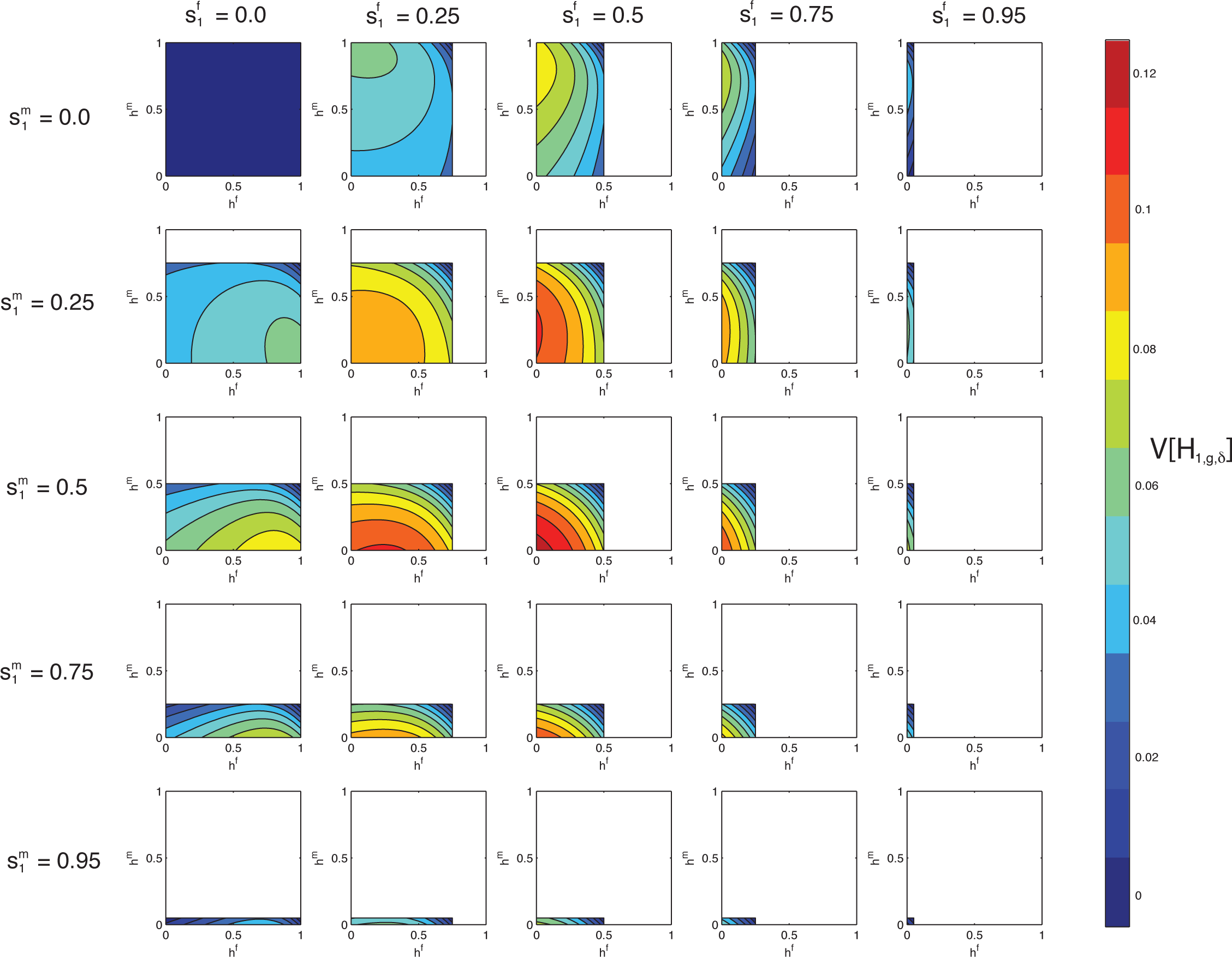
Contour plots of the limit of the variance of the fraction of admixture over time as a function of *h^f^* on the x-axis and *h^m^* on the y-axis for specified values of *s*_1_^*f*^ by column and *s*_1_^*m*^ by row. The domains of *h*^*f*^ and *h*^*m*^ are 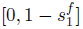 and 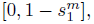 respectively.

As in Figure 9, in the upper left plot in Figure 10, 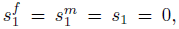 and the limiting variance is a constant zero. In Figure 10, the maximal variance occurs at the origin (*h^f^* = *h^m^* = *h* = 0) of the plot with 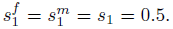 As in Figure 8, at the maximum by eq. (4), *s*_2_^*f*^ = *s*_2_^*m*^ = s_2_ = 0.5. In this case, females and males contribute equally. Both source populations contribute maximally to pull the distribution of the fraction of admixture toward the extremes of zero and one.

Because the limiting variance is symmetrical with respect to source population, and recalling eq. (4), each plot in Figure 10 is equivalent to a corresponding plot in Figure 9 reflected along both the x-axis and y-axis. For example, the plot in Figure 10 with *s*_1_^*f*^ = 0.5 and *s*_1_^*m*^ = 0.25 is equivalent to the Figure 9 plot with 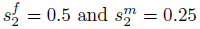 if reflected on both the x-and y-axes.

Figures 8–10 illustrate that the global maximum of the limiting variance occurs when the two source populations contribute equally, the contributions from the two sexes are equal, and the hybrid population does not contribute to the next generation. As the parameters move from the location of the maximal limiting variance to the minimum, the variance monotonically decreases.

### Properties of the limiting variance in terms of 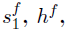 and 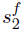

Next we plot on the x-and y-axes two parameters of the same sex from different populations. Because the variance is invariant with respect to transposition of females and males, we consider only females without loss of generality. Specifically, in Figure 11 we plot the limiting variance as a function of *s_1_^f^* on the x-axis and *h^f^* on the y-axis, for fixed values of *s_1_^m^* and *h^m^*. In Figure 12, we plot the limiting variance as a function of *s_1_^f^* and *s_2_^f^* for fixed *s_1_^m^* and *s_2_^m^* For both Figure 11 and Figure 12, the domains of *s_1_^f^*, *s_2_^f^*, and *h^f^* are constrained by eq. (4).

**Figure 11:**
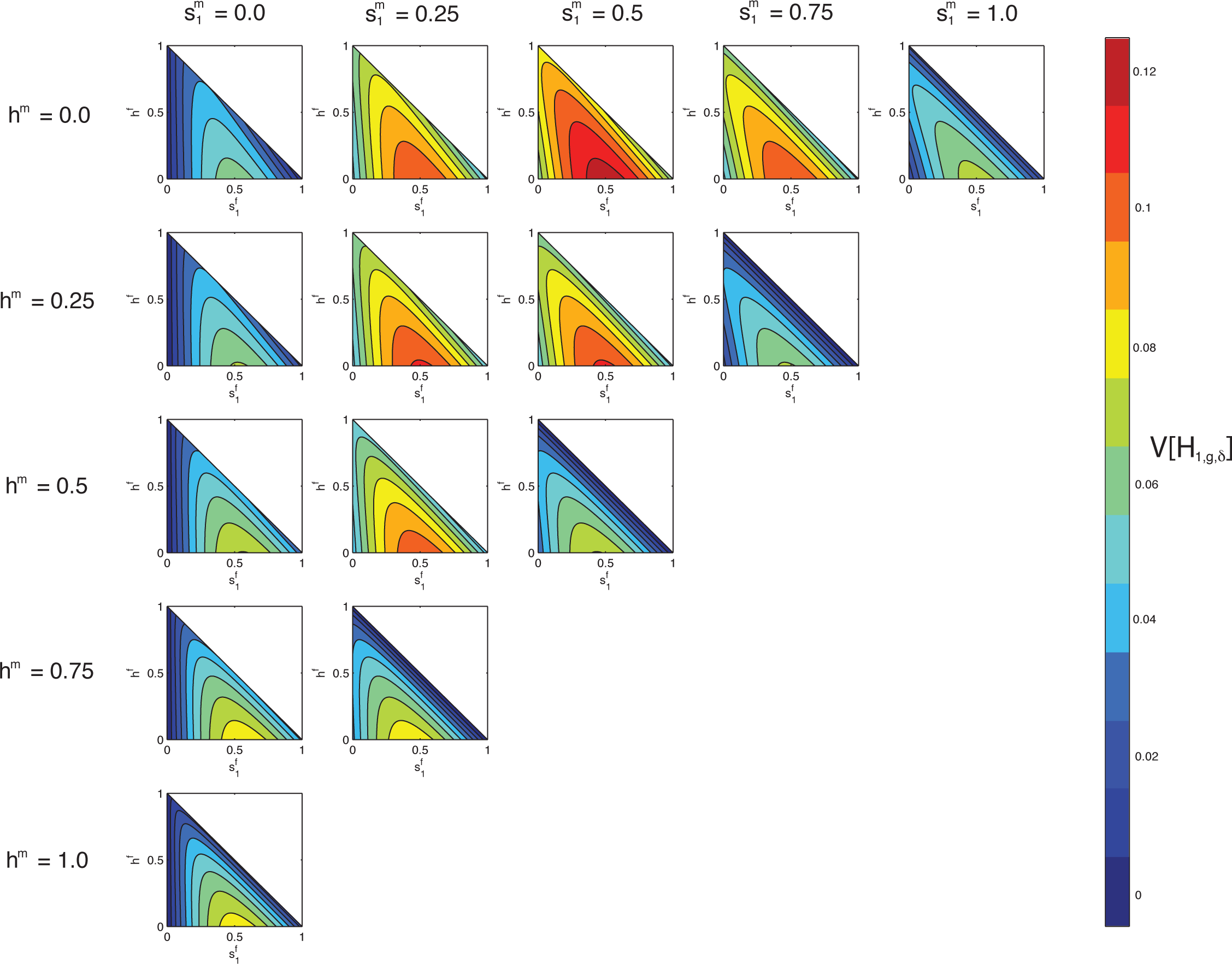
Contour plots of the limit of the variance of the fraction of admixture over time as a function of *s*_1_^*f*^ on the x-axis and *h*^*f*^ on the y-axis for specified values of *s*_1_^*m*^ by column and *h*^*m*^ by row. The domains of *s*_1_^*f*^ and *h*^*f*^ are bounded by the function *s*_1_^*f*^ = *s*^*f*^ = 1.

For Figure 11, the maximal limiting variance occurs in the plot with 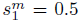 and 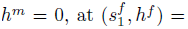 (0.5,0) By eq. (4), this location is the same parameter set for the maximum in Figures 8–10. The maximum in each plot occurs when 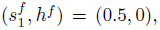 but the magnitude of the variance decreases with increased distance from the plot with fixed *s_1_^m^* = 0.5 and *h^m^* = 0. Similarly, within each plot, the limiting variance decreases with distance from 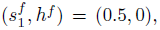.

In the first column of Figure 11, where *s_1_^m^* = 0 the line *s_1_^f^* = 0 produces zero variance because the hybrid population is homogenous, with only one source population contributing. Similarly, on the diagonal *s_1_^m^* + *h^m^* = 1, as *s_2_^f^* = *s_2_^m^* = 0 by eq. (4), the line 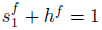 has minimal variance.

In Figure 12, because of the symmetry in sex in eq. (54), the plots above and below those where *s*_1_^*m*^ and *s*_2_^*m*^ are equivalent with a transposition of axes. As in Figures 8-11, the maximal variance occurs in the plot with 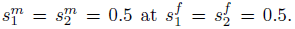 Also, similar to Figure 11, in the first column, when *s*_1_^*m*^ = 0, the line *s*_1_^*f*^ = 0 is a minimum because no contributions trace to source population *S*_1_; in the first row, when *s*_2_^*m*^ = 0 the line *s*_2_^*f*^ = 0 is a minimum.

**Figure 12:**
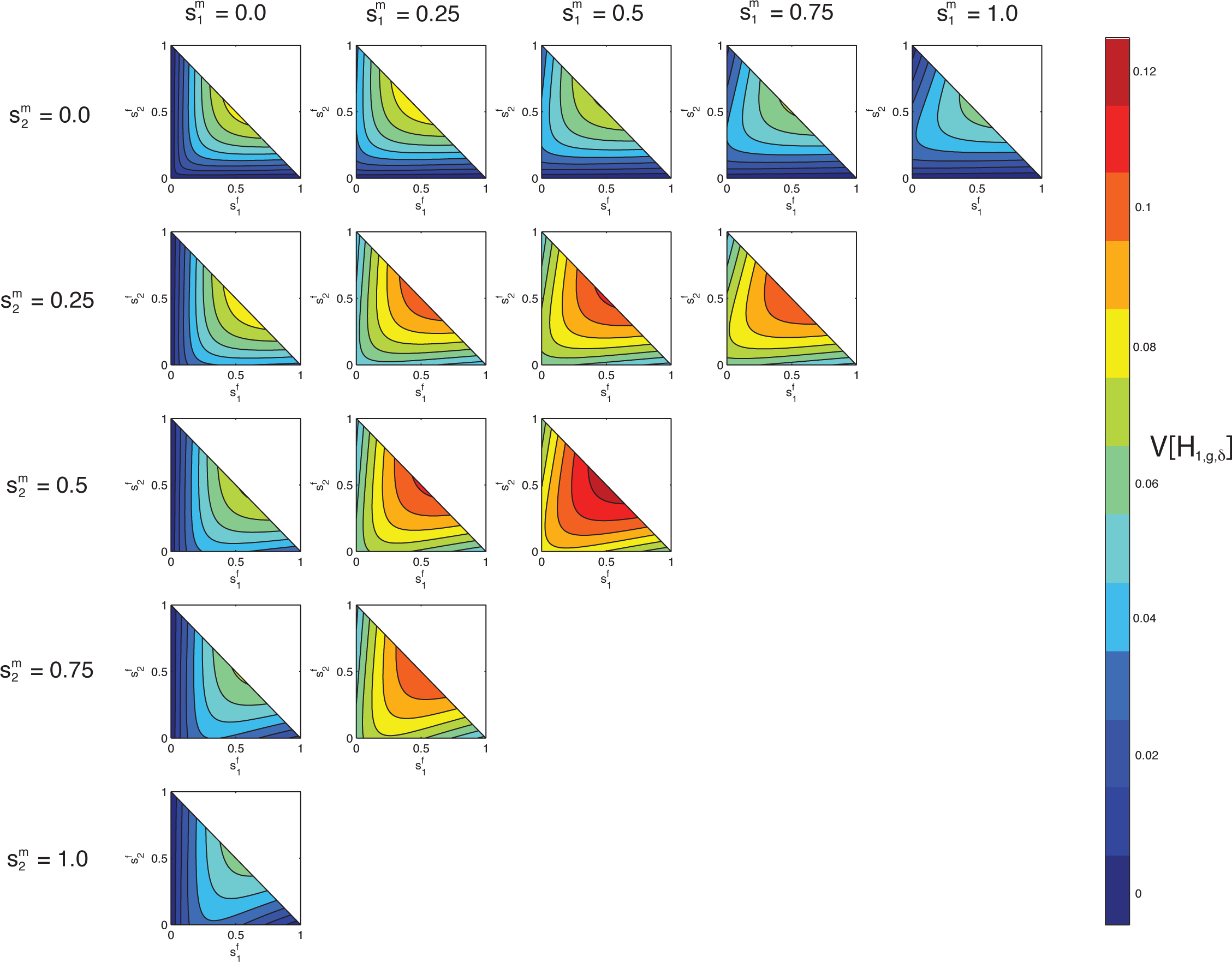
Contour plots of the limit of the variance of the fraction of admixture over time as a function of *s*_1_^*f*^ on the x-axis and *s*_2_^*f*^ on the y-axis for specified values of *s*_1_^*m*^ by column and *s*_2_^*m*^ by row. The domains of *s*_1_^*f*^ and *s*_2_^*f*^ are bounded by the function *s*_1_^*f*^ + *s*_2_^*f*^ = 1

Analogous to the similarity between Figures 9 and 10, by eq. (4), each plot in Figure 12 is a transformation of a plot in Figure 11. For example, for the plot with 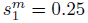 and *h^m^* = 0.5 specified in Figure 11, because the male contributions sum to one, this panel also specifies *s_2_^m^* = 0.25 Therefore, we can compare this plot to the plot with 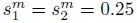 in Figure 12. Both show *s*_1_^*f*^ on the x-axis, and using eq. (4), we can rewrite the y-axis in Figure 12 as *s*_2_^*f*^ = 1−*h^f^*.

### Properties of the limiting variance in terms of non-corresponding parameters

Finally we consider a case in which males from one population in (*S*_1_*, S*_2_*, H*) are compared to females from a different population. While multiple parameter configurations are possible, we plot one that is particularly informative, providing a perspective on eq. (54) beyond the observations visible in Figures 8–12. Figure 13 plots the limit of the variance of the admixture fraction as a function of *s_1_^f^* on the x-axis and *s_2_^m^* on the y-axis, for fixed values of *s*_2_^*f*^ and *s*_1_^*m*^ We rewrite eq. (54) as a function of 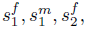 and *s*_2_^*m*^ using eq. (1).

We have,

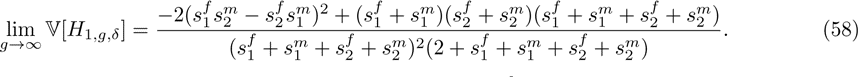

The limit depends on products of sex-specific parameters, including 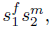 as can be seen in the shape of the contours in Figure 13, but not in the analogous plots in Figure 9.

**Figure 13:**
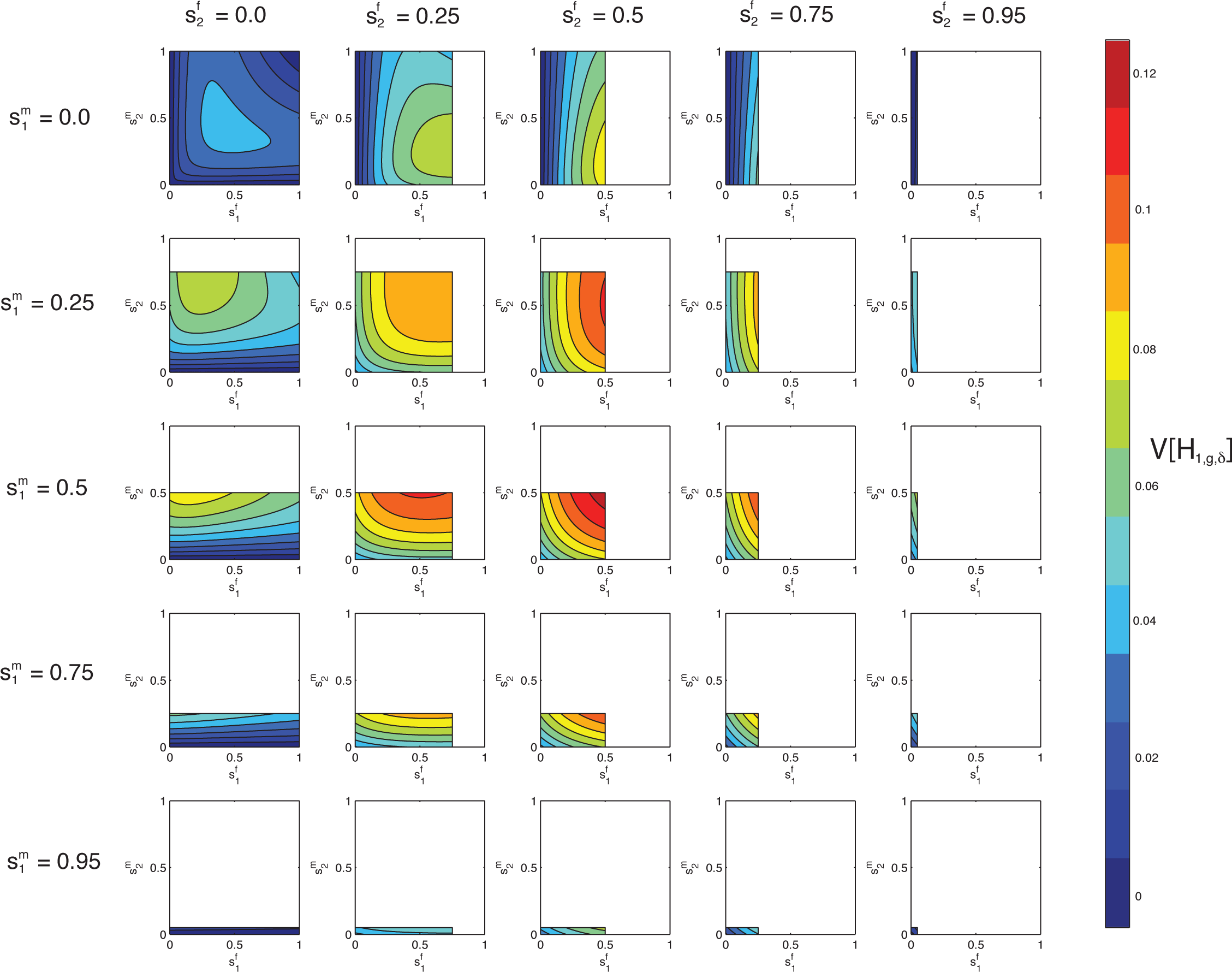
Contour plots of the limit of the variance of the fraction of admixture over time as a function of *s*_1_^*f*^ on the x-axis and *s*_2_^*m*^ on the y-axis for specified values of *s*_2_^*f*^ by column and *s*_1_^*m*^ by row. The domains of *s*_1_^*f*^ and *s*_2_^*m*^ are 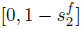 and 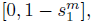 respectively.

## Discussion

Our model demonstrates the potential utility of autosomal DNA in the study of sex-biased admixture histories. Under a framework in which admixture occurs over time, potentially with different male and female contributions from the source populations, we have derived recursive expressions for the expectation, variance, and higher moments of the fraction of autosomal admixture. For the special case of constant admixture over time, we have analyzed the behavior of the variance of the admixture fraction. Although the expectation of the autosomal admixture fraction is dependent only on the total contributions from the source populations, we have found that the variance of the autosomal admixture can be informative about sex-specific contributions. Specifically, for constant admixture over time, we have shown that the variance of the autosomal admixture fraction decreases as the male and female contributions become increasingly unequal.

That autosomal DNA can carry a signature of sex-biased admixture might at first appear counterintuitive, as unlike the sex chromosomes, autosomes are carried equally in both sexes. The phenomenon can, however, be understood by analogy with the well-known result that increasing sex bias decreases the effective size of populations (Wright 1931; Crow & Dennison 1988; Caballero 1994; Hartl & Clark 2007). In a computation of effective size using the coalescent, for example (Nordborg & Krone 2002; Ramachandran et al. 2008), the sex bias causes pairs of genetic lineages to be likely to find common ancestors more recently than in a non-sex-biased population, as the reduced chance of a coalescence in the sex that represents a larger fraction of the breeding population is outweighed by the greater chance of a coalescence in the less populous sex. In a similar manner, if admixture is sex-biased, because lineages are more likely to travel along paths through populations with the larger sex-specific contributions, then the variability of genealogical paths—and hence, the variance of the admixture fraction—is reduced compared to the non-sex-biased case.

Autosomal DNA, with its multitude of independent loci, potentially provides more information about the complex histories of hybrid populations, and the full autosomal genome might be less susceptible to selective pressures at individual loci than the sex chromosomes. To take advantage of autosomal information, many recent efforts to study sex-biased demography have focused on comparing autosomal DNA with the X chromosome (Ramachandran et al. 2004, 2008; Wilkins & Marlowe 2006; Hammer et al. 2008, 2010; Bustamante & Ramachandran 2009; Keinan et al. 2009; Casto et al. 2010; Emery et al. 2010; Keinan & Reich 2010; Labuda et al. 2010; Lambert et al. 2010; Gottipati et al. 2011; Heyer et al. 2012; Arbiza et al. 2014). Our study enhances the set of frameworks available for considering effects of admixture and sex bias on autosomal variation.

For a single admixture event, the expectation of the autosomal admixture fraction is constant in time and not dependent on sex-specific contributions. Unlike in the case of hybrid isolation, if constant nonzero contributions from the source populations occur over time, then the variance of the fraction of autosomal admixture reaches a nonzero limit, dependent on these continuing sex-specific admixture rates, but not on the founding contributions. In both scenarios, the variance can be informative about the magnitude of a sex bias in the admixture history of a hybrid population. For an arbitrary constant total contribution from a source population, the maximal variance occurs when there is no sex bias. The maximal variance across all allowable parameter values of the constant admixture model is seen when there is no sex bias, and equal contributions from both source populations, that is, 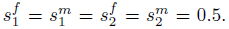 Two admixture histories minimize the variance of the autosomal admixture fraction. First, the variance is zero when only one source population contributes to the hybrid population, and either all hybrid individuals have an admixture fraction of 0 or they all have a fraction of 1. Second, the variance is zero if all males come from one source population and all females come from the other source population. In this scenario, all individuals in the hybrid population have an admixture fraction of 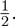

While the variance of the autosomal admixture fraction suggests that autosomal DNA is informative about sex-biased admixture, the relationship between the variance and the sex-specific parameters is complex. We uncovered an interesting case in which quite different sex-specific histories can lead to the same variance over time (Fig. 7). The variance is in fact dependent on the product of multiple sex-specific parameters, but not on each parameter separately (Fig. 13). In particular, when 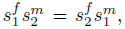 we demonstrate that the variance is maximized (eqs. (54)–(56)). Therefore, when the equality 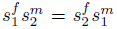 holds, the limiting variance depends only on the total contributions from the source populations, *s*_1_ and *s*_2_ (eq. (57)). The symmetry arises from the non sex-specific inheritance of autosomal DNA.

We have considered two scenarios, isolation of a hybrid population after its founding, and constant contributions from source populations to the hybrid population over time. While the admixture history of real hybrid populations is likely much more complex than these, our models can provide a starting point for statistical frameworks to estimate the parameters of mechanistic admixture models. It is noteworthy that although sex bias does influence autosomal variation, because autosomal DNA is not inherited sex-specifically, the sex that contributes more from a given source population cannot be identified with autosomal DNA alone. Because the X-chromosome follows a sex-specific mode of inheritance, consideration of the X-chromosome alongside autosomal data under the mechanistic model may be able to help differentiate between scenarios that produce the same variance with different choices of the sex with a greater contribution.

## Acknowledgments

We thank Ethan Jewett and Michael D. Edge for useful discussions. We acknowledge support from a National Science Foundation Graduate Research Fellowship and from National Science Foundation grant BCS-1147534.

